# Digital twinning of Cellular Capsule Technology: emerging outcomes from the perspective of porous media mechanics

**DOI:** 10.1101/2020.06.09.142927

**Authors:** Urcun Stéphane, Rohan Pierre-Yves, Skalli Wafa, Nassoy Pierre, Stéphane P.A. Bordas, Sciumè Giuseppe

## Abstract

Spheroids encapsulated within alginate capsules are emerging as suitable *in vitro* tools to investigate the impact of mechanical forces on tumor growth since the internal tumor pressure can be retrieved from the deformation of the capsule. Here we focus on the particular case of Cellular Capsule Technology (CCT).

We show in this contribution that a modeling approach accounting for the triphasic nature of the spheroid (extracellular matrix, tumor cells and interstitial fluid) offers a new perspective of analysis revealing that the pressure retrieved experimentally cannot be interpreted as a direct picture of the pressure sustained by the tumor cells and, as such, cannot therefore be used to quantify the critical pressure which induces stress-induced phenotype switch in tumor cells.

The proposed multiphase reactive poro-mechanical model was cross-validated. Parameter sensitivity analyses on the digital twin revealed that the main parameters determining the encapsulated growth configuration are different from those driving growth in free condition, confirming that radically different phenomena are at play. Results reported in this contribution support the idea that multiphase reactive poro-mechanics is an exceptional theoretical framework to attain an in-depth understanding of CCT experiments, to confirm their hypotheses and to further improve their design.

## Introduction

As a tumor grows, it deforms the surrounding living tissues, which, in turn, produce pressure on the growing tumor and causes strains and associated stresses. Mechano-biology focuses on these mechanical forces and their interplay with biological processes which has been extensively studied experimentally [1–3]. Within this context, current mathematical models of tumor growth are becoming more and more reliable, complement experiments and are useful tools for understanding, explaining and building upon these experimental findings [4–6].

This article focuses on the Cellular Capsule Technology (CCT), an experimental protocol developed by some of the authors in [7] where multi-cellular tumor spheroids (MCTS) were cultured within spherical porous alginate capsules. The latter, after confluence (*i.e.* when the MCTS comes in contact with the inner wall), work as mechanosensors i.e. from their deformation, one can retrieve the stress state within the MCTS. The interaction pressure between the MCTS and the capsule, coming from the basic action-reaction principle, is a capital information since, as envisioned in [7], it could enable the prediction of stress-induced phenotype alterations to characterize cell invasiveness. To this aim, it is essential to quantify the critical pressure which induces the phenotype switch. Notably, it is also relevant to quantify the characteristic time of this process since one can infer that a relatively high pressure sustained by cell during a relatively short time does not lead to phenotype modifications.

Using the measured interaction pressure as a direct discriminant to predict the occurrence of the phenotype switch is very attractive also due to the simplicity of the concept. However, as hypothesized in this paper, directly linking the interaction pressure and the phenotype switch could be a simplistic shortcut since behind the promising practical simplicity of the MCTS-capsule concept, there is a behavior which is not trivially explainable with basic physical concepts. Indeed, the interaction pressure is a quite overall consequence encompassing several mechanisms at a lower level of description. The mechanics of porous media, on which is founded the proposed digital twin of the CCT experiment proposed in this contribution, has emerged as an excellent paradigm to model and possibly reveal these mechanisms offering a new perspective from which one can better interpret and exploit results of the CCT.

The internal structure of a tumor is typically very heterogeneous. Hence, instead of analyzing the system with a homogeneous modeling framework - which is the only option to exploit available data and produce clinically-relevant tumor forecasts (e.g., see [8]), in this contribution the MCTS is modeled as a multiphase continuum consisting of tumor cells, interstitial fluid and an extracellular matrix. This is possible thanks to the richness of experimental data provided by CCT. Mathematical modeling enables retrieving of the stress of each phase from the Biot’s effective stress principle and the adopted multiphase formulation. The model is founded on the rigorous framework provided by the Thermodynamically Constrained Averaging Theory (TCAT) of [9].

To guarantee the scientific relevance of numerical results, the reliability of the model was evaluated using a crossing validation methodology. This has allowed a step-by-step customization of the mathematical model, obtaining a mechanistic formulation which remains predictive also when the experimental conditions of CCT experiment are modified. Systematic sensitivity analyses have been helpful for the analysis and interpretation of results, allowing for quantification of the relative relevance of mechanisms underlying tumor growth phenomenology.

The effective digital twinning of the MCTS-capsule system and emerging biophysical outcomes from the perspective of multiphase porous media mechanics constitute together the novelty of this work. Differently from existing modeling approaches, which are often phenomenological and either too simplistic or too complex that their utility is very weak, the proposed modeling approach is mechanistic and contains the suitable degree of complexity to be representative such a kind of experiment.

## Methods and Model

CCT offers a framework to quantitatively assess the influence of mechanical stresses and its coupling with other biophysical factors impacting tumor cells proliferation and metabolism. Input data for the mathematical model can be retrieved from the CCT experimental conditions; furthermore, numerical results in terms of pressure and displacement can be compared with those measured experimentally. This motivated the selection of CCT as reference experiments.

For the sake of clarity, the experimental observations reported by [7]) together with some additional data provided by the authors are briefly summarized in the following subsection. The mathematical model and the in *silico* reproduction process are then presented.

### Encapsulated MCTS: experimental procedure and observed phenomenology

MCTS cultures have been developed to overcome the limitations of 2D cultures which, inherently, lead to artifacts in cellular behaviors [10], and investigate biophysical aspects. They involve integrin-mediated adhesion, cell differentiation, or drug penetration as a readout for efficient delivery of active species, for which the 3D architecture of the tumor cell aggregate is suspected of having a significant contribution.

MCTS are generally formed and cultured in aqueous medium that we will refer to as “free conditions”. Recently, Alessandri et al. [7] developed a microfluidic technique, namely the CCT, to produce confined MCTS cultures, and they demonstrated that confinement-induced mechanical forces inhibit tumor growth as previously shown [1] but may also trigger a switch towards an invasive phenotype of the tumor cells, with a mechanism that differs from the one mediated by matrix rigidity-sensing [2]. The CCT allows the encapsulation and growth of cells inside permeable, elastic, hollow hydrogel microspheres for the production of size-controlled MCTS. The hydrogel is made of calcium alginate whose pore size of about 20 nm provides a permeability that permits free flow of nutrients and oxygen and ensures favorable conditions for cell proliferation.

All experimental details and useful references can be found in [7], [11]. Briefly, alginate spherical capsules are generated by co-extrusion using a 3D printed device composed of 3 co-axial channels (see figure 1.A): the outermost channel contains the alginate solution; the innermost capillary contains the cells in suspension; the intermediate channel contains an inert (sorbitol) solution that creates a barrier against calcium diffusion from the cell suspension to the alginate solution and subsequent plugging of the device. At flow rates in the 30-45 mL/h range for each channel, a composite liquid jet exits the nozzle and gets fragmented into composite droplets (of radius of the order of the diameter of the nozzle) due to the Rayleigh-Plateau instability. Composite droplets fall into a calcium bath. Since alginate crosslinks almost instantaneously in the presence of divalent ions, hydrogel shells encapsulating cells are readily formed. Then, cellular capsules are washed and transferred to a culture medium and placed in an incubator for temperature and atmosphere control. The capsule therefore serves as a micro-compartment for 3D cell culture.

**Fig 1.**
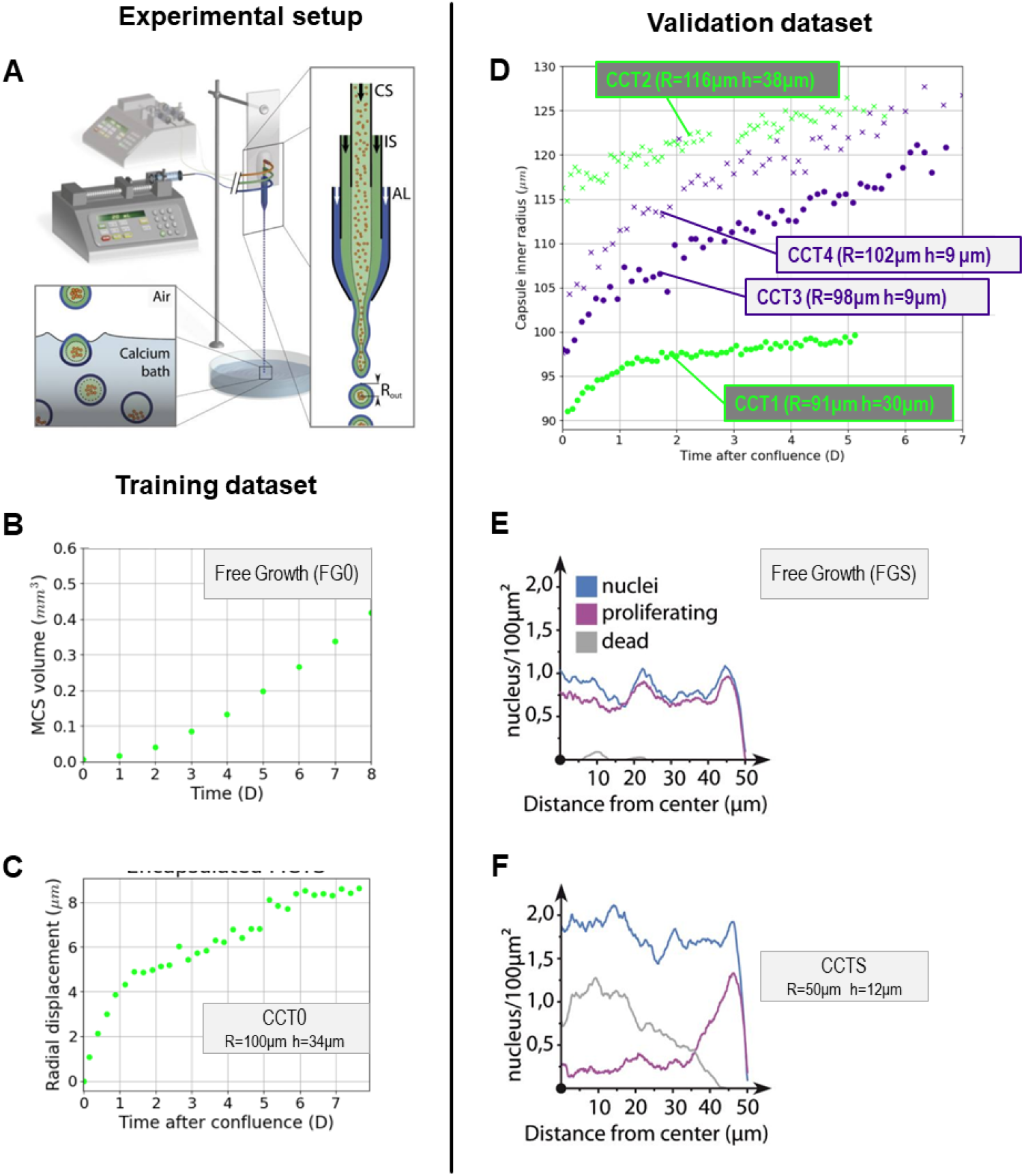
CCT experimental setup and data. **A** CCT microfluidic co-extrusion device; the enlarged view of the chip shows the three-way configuration, with cell suspension (CS), intermediate solution (IS), and alginate solution (AL), respectively, flowing into the coaligned capillaries. **B**-**C** Experimental training data: **B** free growth MCTS control group (FG0), the volume is monitored over a time span of 8 days; **C** encapsulated MCTS, the strain of the capsule (CCT0) is monitored over a time span of 8 days. **D** Validation dataset: two capsules, denoted as thick CCT1 and CCT2; two capsules, denoted as thin CCT3 and CCT4. Their strains are monitored over a time span of 5 to 7 days. **E-F** Experimental quantification of cell nuclei (blue), proliferating cells (purple),and dead cells (gray) along the radius for small free (FGS), **E**, and small confined (CCTS), **F**, spheroids [7].

In the present work, we used CT26 cell lines which are derived from mouse colon carcinoma. During the early stages, cells aggregate with each other. Upon proliferation, the MCTS first grows freely and increases the fraction of the capsule volume occupied by cells until the capsule is filled. By analogy with 2D cell monolayers in a Petri dish, this stage is named confluence. From then on, the MCTS interacts with the alginate shell and deforms it. Conversely, the dilated capsule exerts back a confinement pressure due to action-reaction principle. In this respect, the post-confluent growing MCTS can be regarded as a tumor model that grows against the surrounding tissues and organs. While necrosis at the core of freely growing MCTS due to nutrient diffusion limitation is well known, once the radius of the MCTS exceeds ~ 250-300 *μ*m, similar necrosis can be observed in this experiment, suggesting that both confinement pressure and non-optimal oxygenation due to increased cell density cause these important measurable heterogeneities along the spheroid radius. In order to use the capsules as stress sensors, we need to characterize the material (Young’s modulus *E*_alg_) and morphological (shell thickness, inner and outer radii) properties. First, the dilatation of the alginate capsule was shown to exhibit an elastic deformation with negligible plasticity and no hysteresis. Young’s modulus was measured by atomic force microscopy indentation and osmotic swelling and has been found to be equal to *E*_alg_ = 68 ± 21kPa [7]. Second, and quite remarkably, i) the size of the capsules is set by the size of the nozzle, indicating that different capsule sizes can be obtained by fabricating another microfluidic chip; ii) the thickness of the shell can be tuned with a given chip by varying the ratio between the flow rates in the different channels. For instance, increasing the inner flow rate of the cell suspension with respect to the sum of all flow rates will make the shell thinner. On the basis of these measured parameters, capsules can truly be used as a biophysical dynamometer by using a relation that yields the variation of the inner pressure from the measurement of the capsule radial deformation.

To do so, we used time lapse phase-contrast video-microscopy, which is minimally invasive in terms of photo-toxicity and thus allows monitoring MCTS growth over several days (up to about a week). While MCTS are clearly visible because of the strong light absorption by living tissues, the alginate shell, which is made of 98% water, is fainter. Images were thus analyzed using a custom-made, gradient-based edge detection algorithm that allows tracking simultaneously *R*_MCTS_ and *R*_out_. At time *t* = 0, *R*_in_ was measured manually and did not vary as long as confluence was not reached. In these pre-confluent stages, for all capsule thicknesses, the growth rate of the CT26 spheroid was not significantly different from the one derived from freely growing conditions, indicating that access to nutrients is not compromised by the presence of the alginate shell, whatever its thickness. When confluence was reached (i.e. when *R*_MCTS_=*R*_in_), simultaneous monitoring of *R*_MCTS_ and *R*_out_ allowed computing the variation of the shell thickness, *b*, which was averaged over the capsule perimeter. After confluence, the behavior strongly deviated from that of the free MCTS case. Qualitatively, the same phenomenology was observed for all capsule thicknesses. However, capsule thickness appeard to be a significant determinant of MCTS confined growth when quantitative analysis was performed. Finally, we also reported radial distributions of cell nuclei, dead cells and proliferative cells. These data were obtained i) by fixing the samples at different time points, typically before and after confluence, following standard protocols, then ii) by embedding them in a resin and freezing for cryosection. Immunostaining for nucleus (DAPI) and cell proliferation (KI-67) was performed using two types of fluorophores. Confocal images were then analyzed using standard ImageJ routines for particle detection. To detect dead cells, we exploited the fact that nuclei of dead cells are smaller and brighter (when DAPI-stained) than the ones of living cells. We thus applied additional threshold criteria for brightness and size.

The pressure exerted by the MCTS was calculated using the formalism of thick-walled internally pressurized spherical vessels, and thus could be compared to the stress of each phase from the Biot’s effective stress principle of the multiphase system. Assuming that the alginate gel is isotropic and incompressible the radial displacement of the inner wall, *u*(*R*_in_), reads

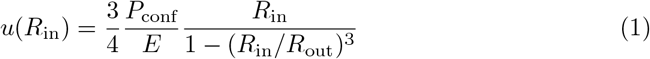

where *P*_conf_ is the internal pressure, *E* is the Young’s modulus, and *R*_in_ and *R*_out_ are the inner and outer radii of the capsule, respectively. Alginate incompressibility also implies volume conservation of the shell. This gives the following constraint equation

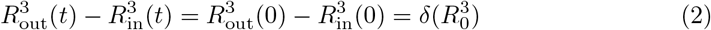

Using this equation, the two time variables *R*_in_(*t*) and *R*_out_(*t*) can be separated and pressure, *P*(*t*), written as a function of *R*_in_(*t*) only

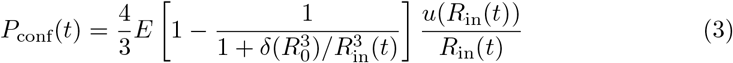

### Experimental input data

First, we considered the denoted *training dataset*:

- for the free MCTS, denoted FG0, the volume was monitored over a time span of 8 days (Fig.1B);
- for the encapsulated MCTS, denoted CCT0, the strain of the capsule (inner radius *R* = 100 *μ*m and thickness *h* = 34 *μ*m) was monitored for 8 days (Fig.1C).

The reliability of the mathematical model was then tested with a *validation dataset* (Fig. 1D):

- Two thick capsules, denoted CCT1 (*R* = 91 *μ*m, *h* = 30 *μ*m) and CCT2 (*R* = 116*μ*m, *h* = 38*μ*m);
- Two thin capsules, denoted CCT3 (*R* = 98 *μ*m, *h* = 9 *μ*m) and CCT4 (*R* = 102*μ*m, *h* = 9*μ*m);

Sparse experimental data have been used to qualitatively measure the model emerging outcomes: the measurements of cell states (quiescent, proliferative, necrotic) of a *R* =50 *μ*m free MCTS denoted FGS (Fig. 1.E) and a small thick capsule, denoted CCTS (*R* = 50*μ*m, *b* =12*μ*m), 26 hours after confluence (Fig. 1.F). Concerning the stiffness of the alginate, *E*_alg_, a range *E*_alg_ = 68 ± 21kPa is provided.

### The mathematical model: a physics-based description of the MCTS-capsule system

Our understanding of the physics and mathematical modeling in oncology has made significant progress owing to our improved ability to measure physical quantities associated with the development and growth of cancer. Hence, health research centers have been collaborating with engineers, mathematicians and physicists to introduce mechano-biology within clinical practice. In [12], the growth inhibition by mechanical stress has been used to reproduce patient specific prostate cancer evolution. This approach can be supplemented by biochemical and genetic approaches, for the prediction of surgical volume for breast cancer [13], or the diffusion of chemical agent in pancreatic cancer [14]. For several years, robust clinical oriented modeling frameworks have emerged (see the seminal article of [8]) wherein multi-parametric MRI patient sets have been processed to initiate patient specific modeling conditions [15]. Recent developments in [16] lead to a deep integration of mathematical oncology in the clinical process, from pre-clinical cell-line growth used for model pre-calibration to multi-parametric MRI for patient specific calibration.

In physics-based modeling, three approaches are currently used to model cancer: discrete, continuum and hybrid (the reader is referred to more detailed descriptions in the work of [17]). Among continuum models, poromechanical ones (e.g., see [6,18–20]) emerge today as valid approaches to model the interplay between biomechanical and biochemical phenomena. As extensively reported in the literature, appropriate elementary models for describing the response of tumour tissue to mechanical and environmental cues will depend on its timescale. At very short time scales (seconds to minutes), tumour cell response is dominated by the elastic response of the cytoskeleton giving tumours a solid-like behavior. For times much longer than a second on the other hand, the response of the cytoplasm to solicitations is essentially viscous and tumour tissue undergoes cellular reorganizations, which lead to large persistent deformations easily represented by a fluid-like viscoelastic model. In this contribution, tumour tissue is modeled neither as a fluid, nor as a solid, but as a multiphasic continuum consisting of a solid matrix (ECM) filled by Newtonian fluid phases.

The application of physics-based models is continuously growing with, for example, *in vivo* modeling reproducing the vascular behavior and experimental validation using histological animal and human samples [5], hybridization of poromechanics and cell population dynamics to mimic the effect of *in vivo* micro-environment [6] or more classic pre-clinical *in vitro* tumor growth [20].

In this paper the multiphase reactive poro-mechanical model of [19] is here further developed and customized for digital twinning of CCT in order to reproduce numerically the experiment of [7] gaining additional information not yet measurable *in vitro*.

Our approach considers the tumor tissue as a reactive porous multiphase system: tissue extra-cellular matrix constitutes the solid scaffold while interstitial fluid (IF) and tumor cells (TC) are modeled as fluid phases. Hence, the mathematical model is governed by momentum and mass conservation equations of phases and species constituting the MCTS-capsule system. Once the capsule is formed, three different spatial domains can be defined (Fig. 2.A): the intra-capsular domain where the tumor cells phase (*t*), the medium/interstitial fluid phase (*l*) and the extra cellular matrix phase (*s*) coexist; the alginate shell domain, where a solid scaffold phase (*s*) and the medium fluid phase (*l*) coexist; and the extra capsular domain where the only medium fluid phase (*l*) exists. In these three domains, strains are calculated according to the theory of poro-elasticity which always assumes the presence of a certain solid phase volume fraction constituting the porous/fibrous medium. Therefore, a certain proportion of the solid phase must always be present even in the extra-capsular domain where it does not exist. Despite this unrealistic condition enforced by the theoretical framework, the reliability of the model is only weakly affected, because the stiffness of this fictitious solid phase is two orders of magnitude lower than that of the alginate solid scaffold (Fig.2.A). A unique physical model is defined for the three domains, with some penalty parameters (*e.g.,* a low intrinsic permeability in the alginate domain) to avoid cell infiltration in the alginate shell. Oxygen advection-diffusion within the medium/interstitial fluid phase is considered. Oxygen acts as the limiting nutrient of TC with prolonged hypoxia leading to the cell necrosis.

**Fig 2.**
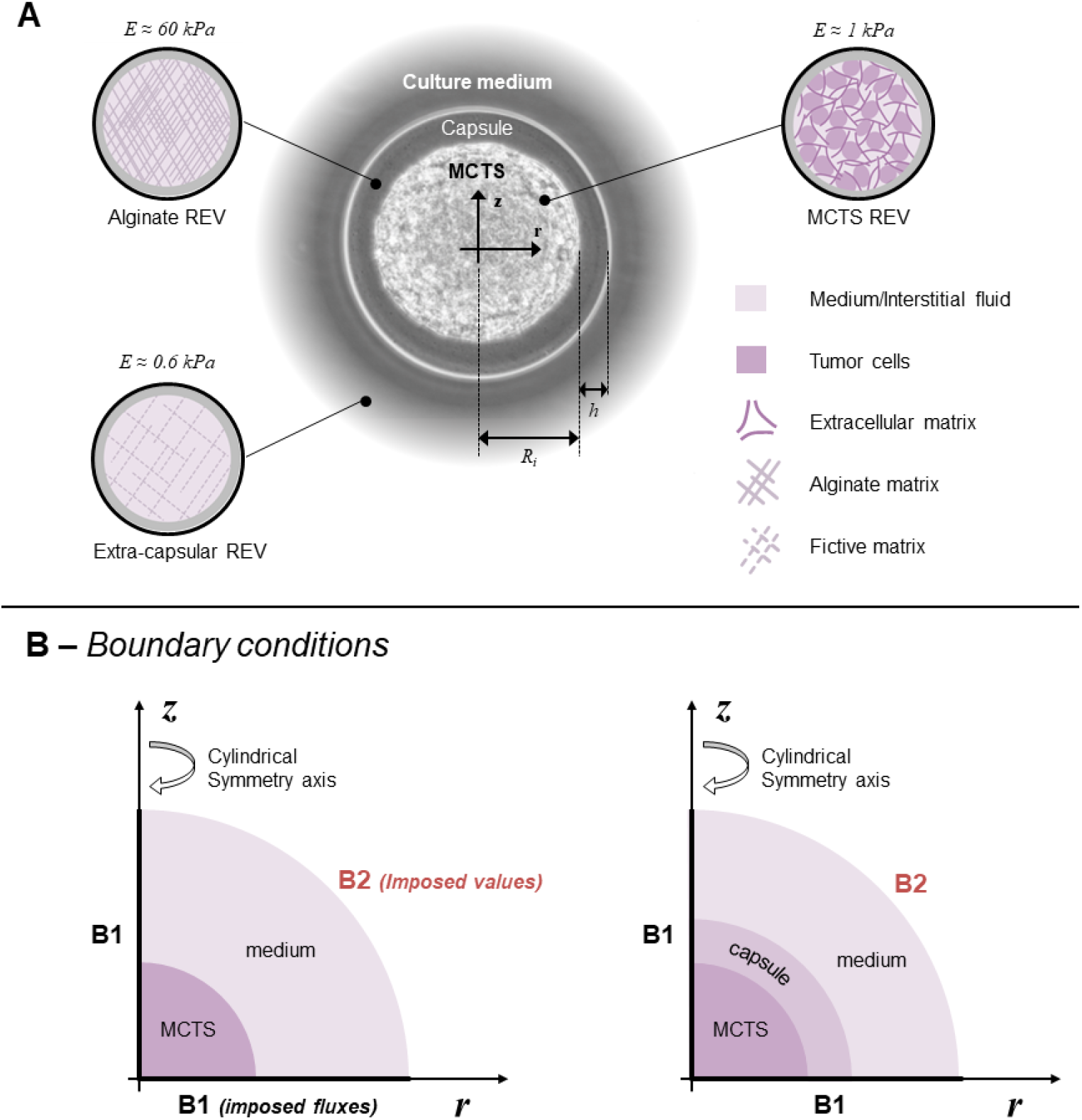
The capsule model. **A** Geometrical description of the capsule and assumed representative elementary volumes (REV): Three spatial domains modeled within the same mathematical framework: MCTS REV (consisting of tumor cells, interstitial fluid (IF) and extracellular matrix), the alginate shell (only IF phase within a solid scaffold of Young Modulus *E*_alg_ = 60kPa) and extra-capsular domain (only IF phase within a fictive solid scaffold of Young Modulus *E*_fict_ = 0.6kPa) enforced by the theoretical framework. **B** Computational boundary condition for the free (left) and confined (right) MCTS. At the bondary B1 symmetry conditions of no-normal flow/displacement are assumed while Dirichlet boundary conditions (e.g., prescribed oxygen concentration) are assumed at the boundary B2

The physical model consists of five governing equations:

- the solid scaffold *s* mass conservation
- the tumor cell phase *t* mass conservation
- the medium/liquid phase *l* mass conservation
- the advection-diffusion equation of oxygen in the medium/liquid phase *l*
- the momentum conservation equation of the multiphase system.

We have four primary variables: three are scalar and one vectorial.

- *p^l^* the pressure of the medium/interstitial fluid
- *p^tl^* the pressure difference between the cell phase *t* and the medium/interstitial fluid *l*
- 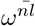 the mass fraction of oxygen
- **u**^*s*^ the displacement of the solid scaffold.

We also have two internal variables: the porosity *ε* and the TC necrotic mass fraction *ω^Nt^.* The evolution of porosity is calculated from the mass conservation equation of the solid phase while the mass fraction of necrotic cells is updated according to the tissue oxygenation in the TC phase (see [18]). We introduce two kinds of closure relationships for the system: mechanical and mechano-biological. Details about the derivation of the governing equations and these constitutive relationships are provided in the following sub-paragraphs.

### The multiphase system

Three phases constitute the multiphase system namely: the solid scaffold *s*, the medium/interstitial fluid *l* and the tumor cells phase *t*. Hence, at each point in the domain, the following constraint must be respected

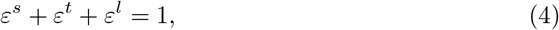

where *ε^α^* is the volume fraction of phase *α*. Defining the porosity *ε* as

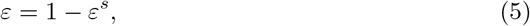

Equation 4 can also be expressed in terms of the saturation degree of fluid phase, *S^f^* = *ε^f^* /*ε* (with *f* = *t, l*)

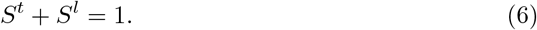

### Mass conservation equations

We express the mass conservation equation for each phase. We use a material description for the motion of the solid phase and a spatial description for the fluid phases, whose reference space is that occupied by the solid scaffold. As the solid is deformable, this reference space is not fixed in time but evolves according to the displacement of the solid phase. For this reason we express mass conservation equations for each phase and species in their material form with respect to the solid scaffold velocity. Mass conservation equations of solid, cell and interstitial fluid phases read:

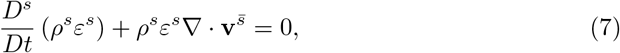

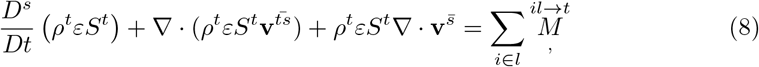

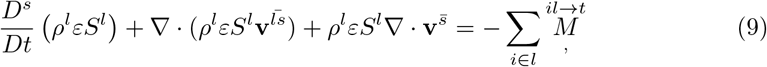

where 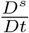 is the material time derivative with respect to the solid phase, *ρ^α^* is the density of phase α, 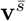 is the velocity vector of the solid phase, 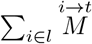 is the total mass exchange (water, oxygen and other nutrients) from the interstitial fluid to the tumor due to cell growth and metabolism, 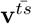 is the relative velocity of cells and 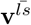 is relative velocity of the interstitial fluid with respect to the solid phase.

The tumor cell phase is a mixture of living (LTC) and necrotic tumor cells (NTC), with mass fraction 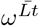 and 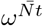, respectively. The following constraint applies

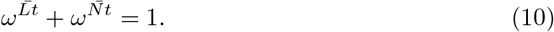

Mass conservation equations for each fraction, assuming that there is no diffusion of both necrotic and living cells, read

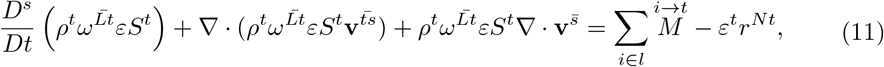

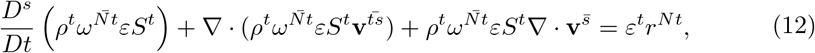

where *ε*^*t*^*r*^*Nt*^ is the death rate of tumor cells. Note that only one of Eqs 11-12 is independent: actually, one can be obtained subtracting the other from Eqn 8 and accounting for the constraint Eqn 10.

Oxygen is the only nutrient which we consider explicitly. Another mass balance equation is introduced which governs the advection-diffusion of oxygen, *n*, within the interstitial fluid

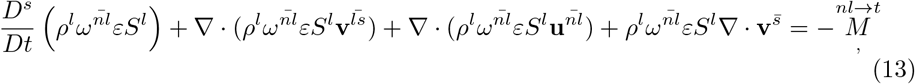

where 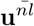 is the diffusive velocity of oxygen in the interstitial fluid and 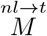 the oxygen consumed by tumor cells due to their metabolism and proliferation rate.

### Momentum conservation equations

We neglect here the effect of gravitational body forces as their contribution is negligible compared to that of other forces. Furthermore, as we assume quasi-static processes and small difference in density between cells and aqueous solutions, inertial forces and the force due to mass exchange can also be neglected. These assumptions simplify the general form of the linear momentum balance equation given in [9] which becomes

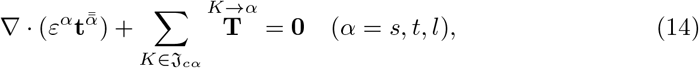

where 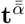 is the stress tensor of phase *α*, 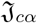 is the set phases connected to *α* and 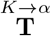 is the interaction force between phase *α* and the adjacent phases. Summing eqn 14 over all phases gives the momentum equation of the whole multiphase system as

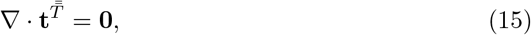

where 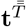 is the total Cauchy stress tensor acting on the multiphase system.

Assuming that for relatively slow flow, the stress tensor for a fluid phase, *f*, can be properly approximated as

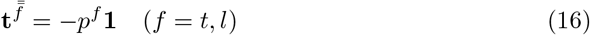

where *p^f^* is the averaged fluid pressure and **1** the unit tensor, Eqn. 14 which apply for a generic phase *α* (solid or fluid) can be expressed in an alternative form for fluid phases as [18]

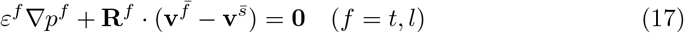

where **R**^*f*^ is a symmetric second-order resistance tensor accounting for interaction between the fluid phase and the solid phase, *s*. Eqn. 17 can be rewritten as

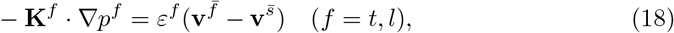

where **K**^*f*^ = (*ε^f^*)^2^(**R**^*f*^)^−1^ is called the hydraulic conductivity. The hydraulic conductivity depends on the dynamic viscosity of the flowing fluid, *μ^f^*, on the intrinsic permeability of the porous scaffold, *k*, and on the fluid saturation degree, *S^f^*, via a relative permeability function 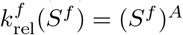 (*A* depending on the fluid characteristics, see [21,22]). As customary in biphasic flow problems we set here 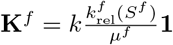. Hence, the governing linear momentum conservation equations for tumor cells and interstitial fluid read respectively

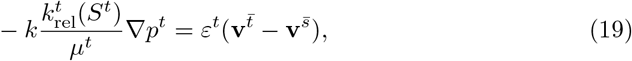

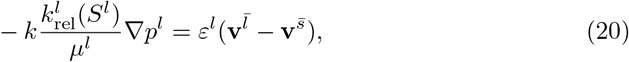

### Effective stress principle

We assume here that all phases are incompressible. However, the overall multiphase system is not incompressible, because of the presence of porosity that evolves according to the scaffold deformation. As all phases are incompressible, their densities *ρ^α^* (with *α* = *s, t, l*) are constant and the Biot’s coefficient is equal to 1. With these premises, the total Cauchy stress tensor appearing in eqn 15 is related to the Biot’s effective stress as follows

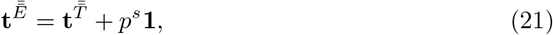

where *p^s^* = *S^t^p^t^* + *S^l^p^l^* is the so-called solid pressure, describing the interaction between the two fluids and the solid scaffold.

The chosen closure relationship for the effective stress 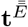 is linear elastic:

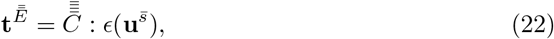

with 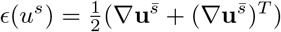 and 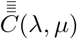 the fourth order elasticity tensor, reduced in Voigt notation: 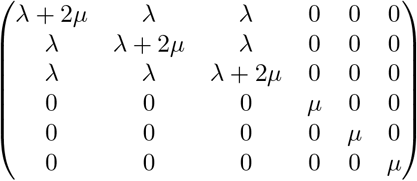 with the Lamé constant 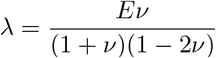 and 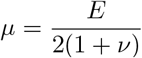.

*E* is the Young modulus of the solid scaffold and *ν* its Poisson ratio.

### Pressure-saturation relationship

The experimental measurement of cells density inside the capsule revealed a strong dependency to necrotic fraction 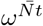. Hence, the pressure-saturation closure relationship has been improved with respect to that proposed in [19], to be more physically relevant and adapted to confinement situation

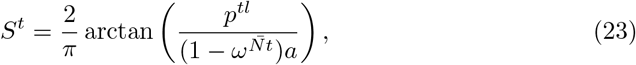

with *p^tl^* pressure difference between tumor and interstitial fluid (i.e. *p^tl^* = *p^t^* – *p^l^*). The saturation is directly linked to the partial pressure of the phase and a constant parameter *a*, which accounts for the effect of cell surface tension and of the refinement of the porous network (see [22] for the biophysical justification of the proposed equation). Its influence is offset by the necrotic fraction of tumor cells, 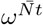 (see Fig. 3), which allows us modeling necrotic areas of very high cell density according to experimental evidence.

**Fig 3.**
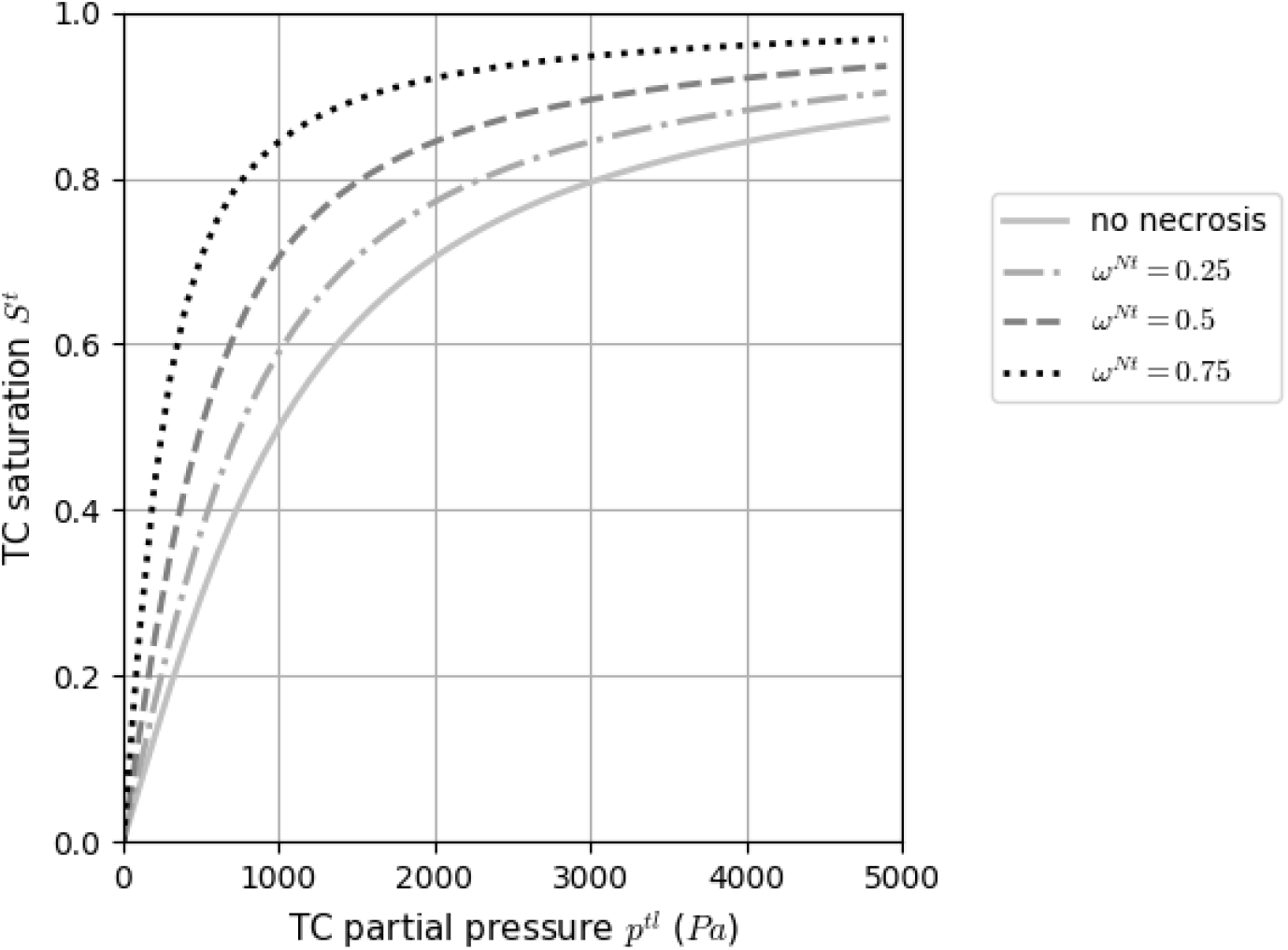
Constitutive relationship between tumor cell partial pressure and saturation. Tumor cell phase saturation *S^t^*, with the parameter *a* (fixed to 1kPa in the figure), evolving with the necrotic fraction of the phase 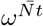

### Nutrient diffusion

The tumor cells growth, metabolism and necrosis are regulated by a variety of nutrient species and intracellular signalling. However, without losing generality, in the present model one single nutrient is considered: oxygen. The case of multiple species can be easily obtained as a straightforward extension of the current formulation. The Fick’s law, adapted to a porous medium, was adopted to model diffusive flow of oxygen eq.13:

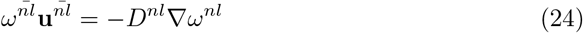

where *D^nl^* the diffusion coefficient for oxygen in the interstitial fluid is defined by the constitutive equation from [21]

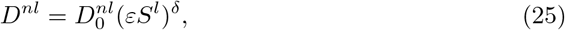

the exponent *δ* it set equal to 2 to account for the tortuosity of cell-cell interstitium where oxygen diffuse ([19], [20], [23]).

### Tumor cells growth, metabolism and necrosis

Tumor cell growth is related to the exchange of nutrients between the IF and the living fraction of the tumor. The total mass exchange from IF to the tumor cell phase is defined as

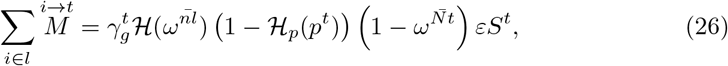

Note that 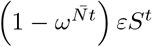 is the living fraction of the tumor. 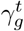 is the tumor growth rate parameter, cell-line dependent. 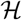 and 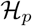 are regularized step functions varying between 0 and 1, with two threshold parameters *σ*_1_, *σ*_2_, that is to say 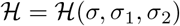. When the variable *σ* is greater than *σ*_2_, 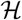 is equal to 1, it decreases progressively when the variable is between *σ*_1_ and *σ*_2_ and is equal to zero when the variable is lower than *σ*_1_. 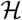 represents the growth dependency to oxygen:

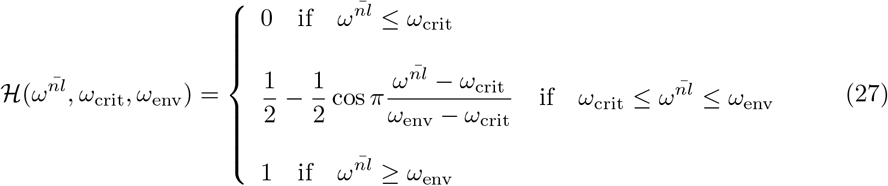

*ω*_env_, the optimal oxygen mass fraction, is set to 4.2 * 10^−6^ which corresponds, according to Henry’s law, to 90mmHg, the usual oxygen mass fraction in arteries (see [24]). *ω*_crit_, the hypoxia threshold, is cell-line dependent, for tumor cells, it has been set to a very low value: 10^−6^ (≈ 20mmHg, for common human tissue cells, hypoxic level is defined between 10 and 20mmHg [25]) The function 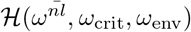 is plotted Fig. 4A.

**Fig 4.**
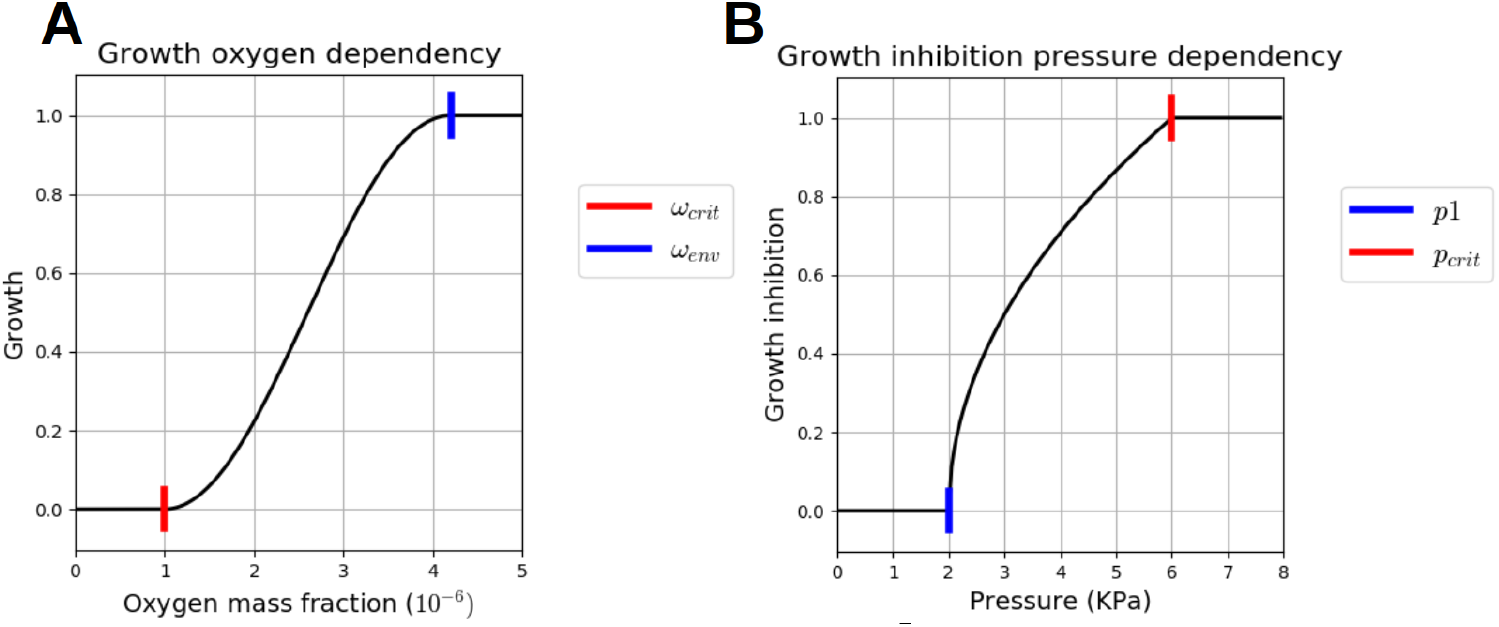
Two mechano-biological laws. **A** 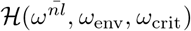. The TC growth and nutrient consumption are dependent to the oxygen mass fraction 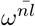. If it is lower than *ω*_crit_, the TC growth is stopped and the nutrient consumption is reduced to the metabolism needs only. If it is greater or equal to *ω*_env_, the growth and the nutrient consumption are maximum. **B** 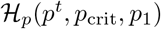. The TC growth and nutrient consumption are dependent to the TC pressure. If it is greater than *p*_1_, the 2 processes begin to be strongly affected and if the TC pressure reaches *p*_crit_, they are totally stopped.

Function 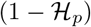 represents the dependency on pressure:

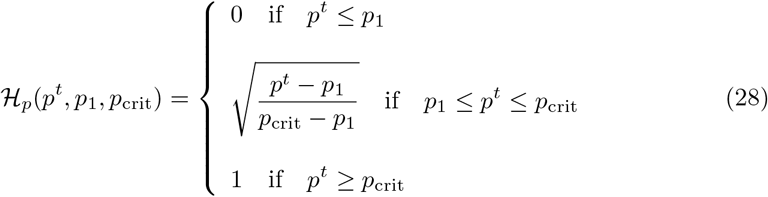

An example of the function 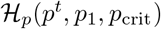 is plotted Fig. 4B, we have set *p*_crit_ to 6 kPa as initial guess (in [1], they found a inhibitory pressure at 10 kPa) and *p*_1_, the pressure threshold when the inhibitory process starts, at 2 kPa.

As tumor grows, nutrients are taken up from the IF so that the sink term in eq.13 takes the following form:

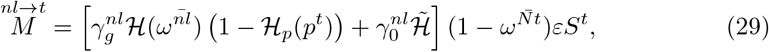

Nutrient consumption from IF is due to two contributions: the growth of the tumor cells, as given by the first term within the square brackets in eq.29, the metabolism of the healthy cells, as presented in the second term. Thus, 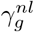 is related to the cell proliferation, as discussed above; whereas the coefficient 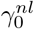 relates to the cell metabolism. 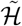 is an adaptation of the previous step functions for the cell metabolism:

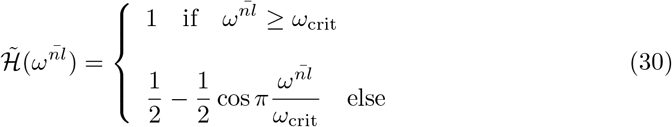

The model does not discriminate between proliferating and quiescent cells, but the growth is subject to 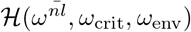. To make possible the comparison with the experimental proliferative cell quantities (see Fig. 9), the following relationship has been set:

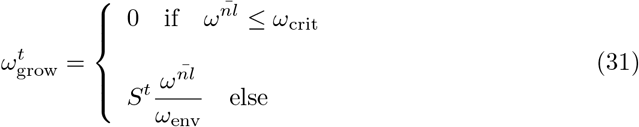

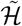 is also used in the definition of hypoxic necrosis rate which reads

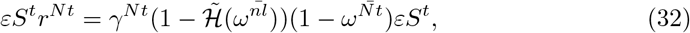

where *γ^Nt^* = 0.01 is the necrotic growth rate. As the experimental data on necrosis were too sparse for this parameter identification (only a few stained-cell imaging), we have kept its generic value.

### Initial parameter settings

As prescribed in [26], aiming for biological or clinical relevancy demands to investigate the choice of the initial values for each of the parameters. Some parameters are of physical nature (the IF dynamic viscosity, the oxygen mass fraction inside cell cultures), they can be, sometimes with enormous efforts, measured (or at least their values will be compared to the physical soundness). Others parameters belong more specifically to bio-poromechanical models in the mathematical oncology fields. Some of them have quite a theoretical nature (*e.g.,* the ’permeability’ of the ECM) while others have been experimentally measured at the cellular level (*e.g.,* the oxygen consumption rate of EMT6/Ro cell line in [27]). For these parameters, we have taken values that previous numerical studies ([28], [19], [20], [23]) have used for MCTS cultured with other cell lines (human glioblastoma multiforme and human malignant melanocytes), averaged these values, denoted *’generic’,* and used them as initial guess for identification of parameters of our CT26 cell line based MCTS.

When experimental data did not provide any relevant information on a parameter (e.g. for ECM stiffness and permeability), we chose to fix them at their generic value. The following parameters have a non negligible influence on the model outputs, and the closure relationships they belong to are explained in detail in the mathematical model section: 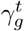 the TC growth rate (Eq.26), 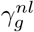 and 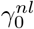 the oxygen consumption rate due to growth and quiescent metabolism respectively (Eq.29), *a* the parameter tuning the joint impact of the ECM thinness and cell surface tension (Eq.23) and *μ_t_* the TC dynamic viscosity, presented in Eq.19. Two other parameters, *p*_1_ and *p*_crit_, are introduced in this modeling. They represent thresholds which govern the inhibition of the proliferation (Eq.28) of cancer cells. The initial guess of *p*_crit_ have been chosen according to the work of [1] and [2], and the initial guess of *p*_1_ has been set by observation of the experimental data.

### In silico reproduction process

From the computational point of view, we aimed to a light and adaptable process: free, open source and compatible with any 2D or 3D geometry. For the model validation, we followed the convention of mathematical oncology proposed in [26]: two distinct sets of data for optimization and validation, the parameters set being fixed before validation. To measure the quality of the fits, we followed the prescription of [29]: the root mean square error (RMSE) relative to a reference, specified each time. The error on the numerical quantity *ξ*_num_ relative to a reference *ξ*_ex_, evaluated at *n* points is computed as:

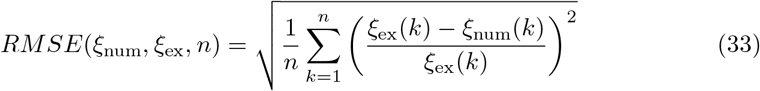

#### Computational framework

We implemented the above model in Python and C++ within the FEniCS framework [30], with an incremental monolithic resolution of the mixed finite element (FE) formulation. The simulations have been run with composite Taylor-Hood element *P*_3_(ℝ^2^), [*P*_2_(ℝ)]^3^ (one vectorial and three scalar unknowns), a mesh cell size of *dh* = 5 *μ*m and an implicit Euler scheme with *dt* = 1200 s. All the details and analytical verification of the FE formulation can be found in Appendix A. Computational framework. All the codes used in this article, analytical verification, integration along the inner radius of the radius, free growth and confined growth, are available on Github, at https://github.com/StephaneUrcun/MCTS_mechanics

#### Initial and boundary conditions

The Fig. 2 shows the two modeled configurations of MCTS (the free on the left and the confined on the right). Each mesh is half of a sphere because we also exploit symmetry with respect to a diametrical plane. For the three scalar variables, we prescribed Dirichlet boundary conditions along the outer radius of the domain *p^l^* = 0, *p^tl^* = 0 and 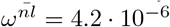 (which corresponds, according to Henry’s law, to 90 mmHg, the usual oxygen mass fraction in arteries, see [24]) and no flux condition at *r* = 0 and *z* = 0. For the ECM displacement field **u**^*s*^, slip conditions 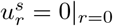 and 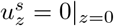 are used, and Dirichlet conditions **u**^*s*^ = **0** at the outer radius of the domain (see Fig. 2).

#### Local sensitivity analysis

We performed a local sensitivity analysis to estimate Sobol sensitivity indices on the FG0 and CCT0 training datasets to assess the sensitivity of the FE solution to the input parameters, both on the free and encapsulated MCTS. Further details of the process can be found in Appendix B. Sensitivity analysis. First, we designed two cost functions, for FG0 and CCT0. The free growth cost function, *J*_free_, compares the experimental aggregate volume *V*_exp_ and the simulated volume from day one to day four.

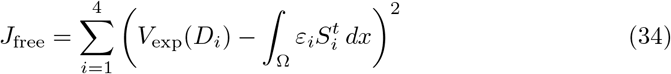

The cost function for CCT0, *J*_conf_, compares two experimental quantities, the displacement of the internal radius **u**(*R*_in_) and the internal pressure calculated in Eq.3, with the corresponding model outputs **u**^*s*^ and *p*^*s*^, one day after confluence.

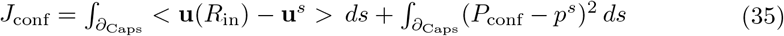

where *∂*_Capsule_ means along the inner radius of the capsule.

The results of the two configurations, with the parameters at their initial values used for the sensitivity analysis can be found Fig.14 Appendix B. Sensitivity analysis. Secondly, the 7 parameters 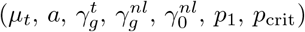 were disturbed one at a time respectively to a [–10, –5, –2, –1, +1, +2, +5, +10]% grid. The variations of *J*_free_ and *J*_conf_ were interpolated by a linear model, which constitute first-order Sobol indices. The influence of the *i^th^* parameter 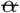 was deduced from the slope 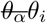 of the linear fit. The Sobol index 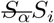 was calculated as follows:

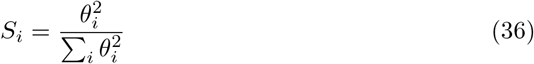

Finally, the 21 parameters tuples were evaluated at the 2 extreme values of the grid, [–10, +10]%, for each configuration. The variations of the solution *J*_free_ and *J*_conf_ were interpolated by a second-order polynomial model. This allowed computing two types of Sobol indices: *S_i_* for the influence of the parameter *i* and *S_j_* for the influence of each couple (*i,j*) of parameters.

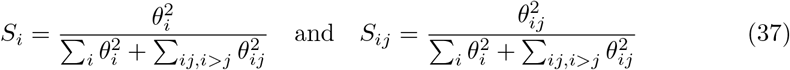

#### Parameter identification and model validation

For both configurations, the optimization procedure was based on sensitivity profiles, that is to say, we identified the set of parameters that gathered at least 90% of the variance of the solution. Then, the selected parameters were identified by a Nelder-Mead simplex algorithm (in the Python library SciPy, method minimize, option Nelder-Mead). In one hand, this algorithm has the advantage of not requiring the computation of the system gradient to the parameters, and, on the other hand, generally converges to a local minimum. To avoid this phenomenon, a large range of initial guess was tested: [–20, +20]% around the *generic* values for 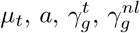, and 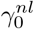 and [–50, +50]% around the initial guess of *p*_1_ and *p*_crit_, as we have not previous literature values. All the parameters having a physical meaning, their values were bound to the physical and physiological values reported in the literature, and we choose not to extend this range. In the CCT0 configuration, considered as the representative case by the team of [7], the parameters of the MCTS cell-line were identified with the mean experimental value of the alginate stiffness: *E*_alg_ = 68 kPa; it is important to note that this study is not aiming to identify the stiffness of this biomaterial.

To evaluate the reliability of the identified parameters, an author of this article and member of the team of [7] have jointly provided unpublished experimental results of encapsulated MCTS, CCT1 to 4 and CCTS, namely the validation dataset. As their alginate stiffness is not known (the Young’s modulus of the alginate is estimated to be *E*_alg_ = 68 ± 21 kPa), two simulations were run for each capsule with the extreme values of Ealg. This provided the range of modeling possibilities of the identified parameters (Fig 6, grey range), for each capsule an indicative fit is proposed with a value of *E*_alg_ which minimize the RMSE.

## Results

Based on a detailed sensitivity study, the identified set of parameters was tested and cross-validated on unpublished experimental results (Fig. 1.C-E) provided by the same team of [7]. Numerical simulation also provides a wide output of qualitative results which are presented and interpreted. At the end of the section, we show that the model outputs allow predicting, with a reasonably good accuracy, experimental TC saturation and its necrotic fraction, despite these quantities have not been used for the model optimization.

### Sensitivity analysis

Fig. 5 shows the results of first-order and second-order interaction analyses, for the free and encapsulated configurations respectively. Clearly distinct profiles were obtained.

**Fig 5.**
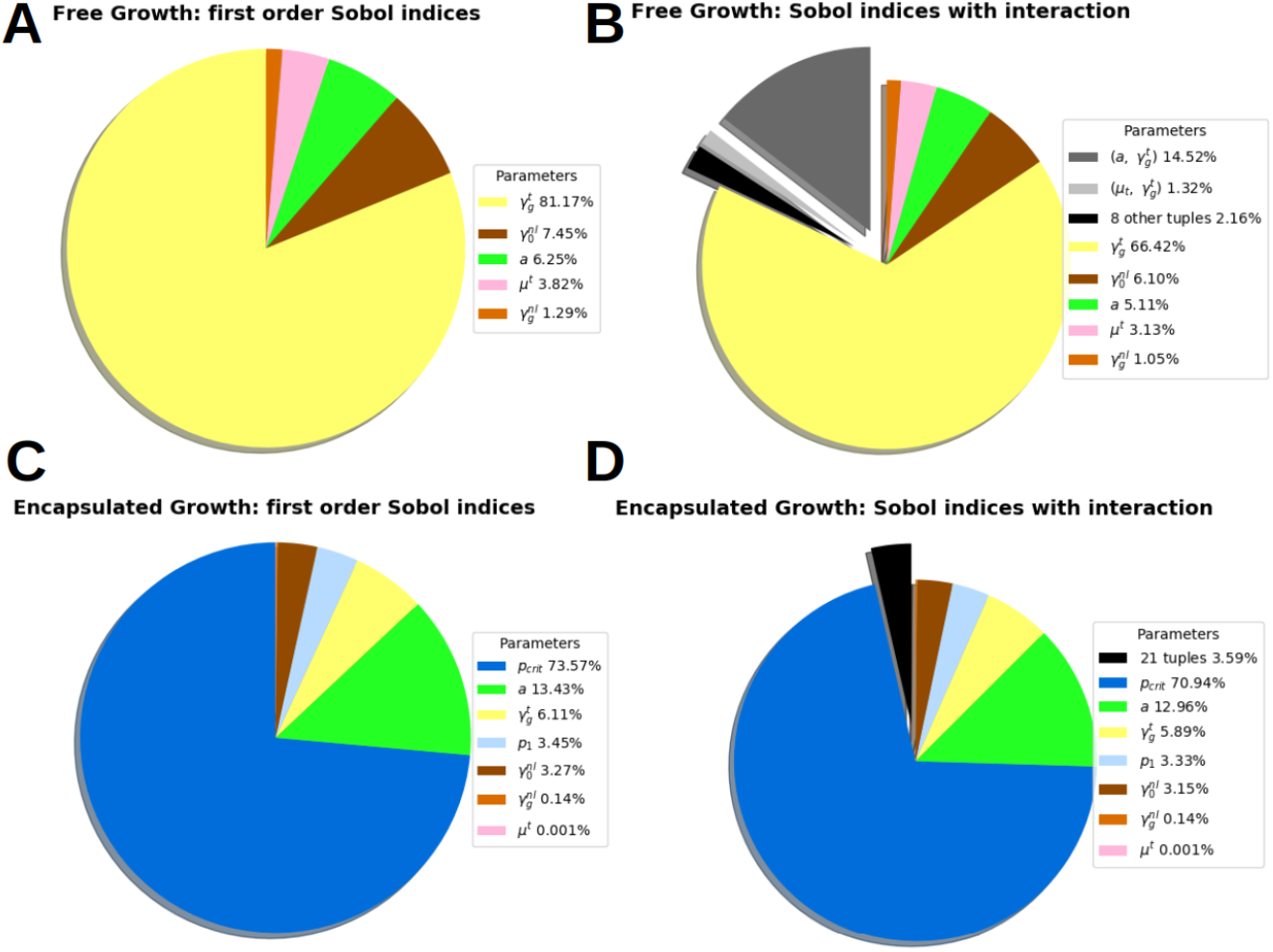
Sobol indices of the solution sensitivity. Sobol indices of the solution sensitivity on 7 parameters: *μ_t_* the TC dynamic viscosity, the parameter *a* accounting for the joint impact of the ECM thinness and cell surface tension, 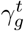 the TC growth rate, 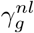 and 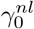 the oxygen consumption rate due to growth and quiescent metabolism respectively, *p*_1_ and *p*_crit_, two thresholds which govern the pressure-induced inhibition of the TC proliferation. Free MCTS configuration (top row). **A First-order analysis**: Only 5 parameters remain, the governing parameter is 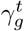, the tumor cells growth rate, the sensitivity of the solution on the pressure parameters, *p*_1_ and *p*_crit_, is 0. **B Interaction**: among 10 parameters tuples, one is significant 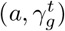 accounting for 14.5% of the solution variance. Thus, these two parameters are not independent and should identified together. The total variance of the solution shows that, considering all the interactions, the influence of each parameter alone is not qualitatively changed: 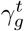 from 81.1% to 66.4%, 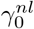 from 7.4% to 6.1%, a from 6.2% to 5.1%. Encapsulated MCTS configuration (bottom row). **C First-order analysis**: the governing parameter is *p*_crit_ the inhibitory pressure of tumor cells growth (73.5% of the solution variance). **D Interaction**: the sum of 21 parameters tuples represents 3.6% of the solution variance (the detail of 21 tuples can be found in Appendix B. Sensitivity analysis, table 7).

In the free growth control group FG0, the governing parameter is 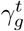 the tumor cells growth rate (first-order index, 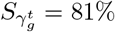, with interactions 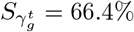). In decreasing order of magnitude, we observed that two parameters are not negligible: 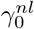 the oxygen consumption due to metabolism (first-order index 7.45%, with interactions 6.10%) and *a*, the parameter determining the joint impact of the ECM thinness and cell surface tension (first-order index 6.25%, with interactions 5.11%). The important difference between the 2 Sobol indices of 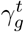 is explained by the only non negligible interaction between two parameters: 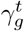 and 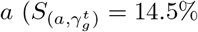, see Fig. 5, right). This important interaction is indicative of the significant roles that ECM properties and cell-cell adhesion have on proliferative-migration behavior (this is widely described in literature, see for instance [2]) and that our modeling approach can reproduce mechanistically how these properties impact the overall observed phenomenology of tumor growth. Thus, these two parameters are not independent and should be identified together.

For all parameter perturbations in the first-order and second-order interaction analyses, the pressure of the TC phase *p^t^* = *p^l^* + *p^tl^* was less than 1 KPa, thus the first threshold of growth inhibitory due to pressure *p*_1_ was never reached and, *a fortiori,* the critical threshold of total inhibition *p*_crit_. Thus, the sensitivity of the FE solution to *p*_1_ and *p*_crit_ was 0. The 3 parameters 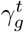, *a* and 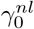 has therefore been optimized for the free configuration.

For the encapsulated configuration CCT0, the governing parameter is the critical inhibitory pressure *p*_crit_ (first-order *S*_*p*_crit__ = 73.5%, with interaction 70.9%). 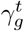, *a* and 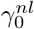 has already been identified for the free configuration, the only non negligible parameter remaining is *p*_1_ (first-order index *S*_*p*1_ = 3.4%, with interaction 3.3%). The difference between Sobol indices of first-order and interactions is weak. Indeed, the 21 parameters tuples capture only 3.6% of the solution variance. Thus, in the encapsulated configuration the parameters can be considered non-correlated and be identified separately.

Such results allow us highlighting that, in the encapsulated configuration CCT0, the mechanical constraint is the phenomenon that determines the overall growth phenomenology provided by the mathematical model.

### Calibration

The three governing parameters 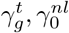, *a* for the free MCTS configuration were identified using the Nelder-Mead simplex algorithm and fitted to the experimental data with a *RMSE* = 0.031. To be physically relevant, the same parameters set should be shared by the two configurations. These three parameters being calibrated, we consider they describe a limited part of the mechano-biological states and they are thereafter fixed and injected in the encapsulated configuration. Its two remaining parameters *p*_1_, *p*_crit_ (74% of the variance) were identified using the same algorithm. We fitted the experimental data of the encapsulation with a *RMSE* = 0.124. Fig. 6A shows the two configurations fitted with the following set of parameters: 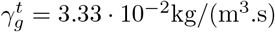, 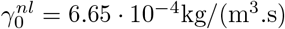, *a* = 890 Pa, p_1_ = 1432 Pa, *p*_crit_ = 5944 Pa (see Table 1). This set is cell-line specific, only relevant for CT26 mouse colon carcinoma.

**Fig 6.**
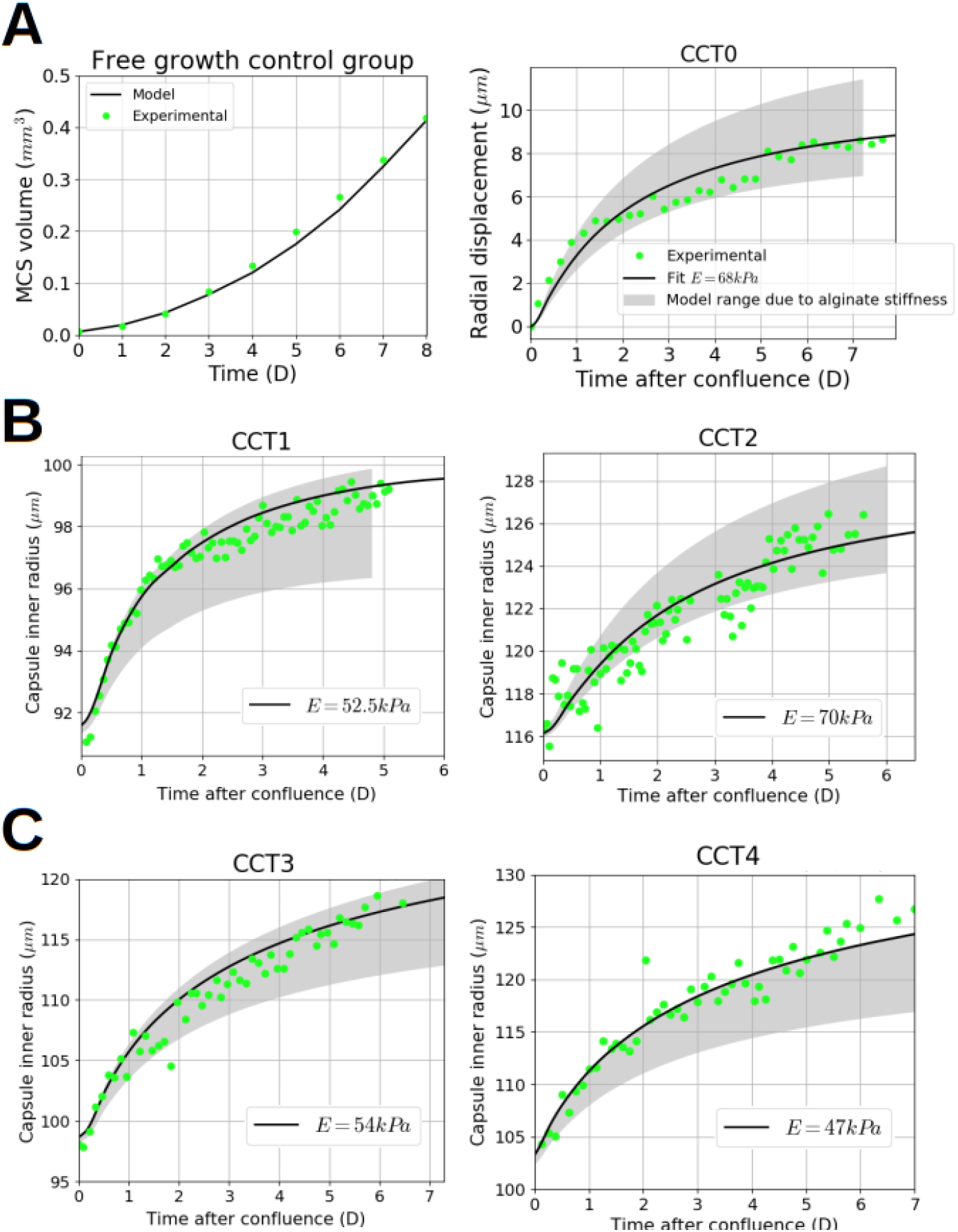
Validation of the calibrated parameters. Alginate Young’s modulus was estimated in [7] as *E*_alg_ = 68 ± 21 kPa. Simulations with the extreme values of *E*_alg_ give the range of possibilities of the optimized set predicted with the model. Experimental results, green dotted; Numerical results with the optimized parameters set, black; Modeling range, grey filled. **A**. Left, free MCTS control group, Time (Day) versus MCTS volume (mm^3^), the model fits the experimental data with a *RMSE* = 0.031. Right, CCT0. The fit uses the mean experimental value of the alginate stiffness *E*_alg_ = 68 kPa. The model fits the experimental data with a *RMSE* = 0.124. **B** Validation of the identified parameters on 2 thick capsules, CCT1 and CCT2. Time (Day) versus Capsule radius (*μ*m). The experimental points are in the modeling range. Both capsule fit with *E*_alg_ = 52.5 kPa and *E*_alg_ = 70 kPa respectively. **C** Validation of the identified parameters on 2 thin capsules, CCT3 and CCT4.. Time (Day) versus Capsule radius (*μ*m). Left, one capsule is fitted with *E*_alg_ = 54 kPa; right, an important part the experimental points are outside of the modeling range.

**Table 1.** Parameters for the CT26 cell line. Source of the generic values: [28], [19], [20], [23].

| Parameter | Symb. | Generic | Unit | Optimized |
| --- | --- | --- | --- | --- |
| ECM network thinness | $a$ | 800 | Pa | 890 |
| Dynamic viscosity of TC | $\mu_t$ | 36 | Pa.s | negligible |
| TC growth rate | $\gamma_g^t$ | $4 \cdot 10^{-2}$ | kg/(m<sup>3</sup>.s) | $3.33 \cdot 10^{-2}$ |
| TC growth $O_2$ consump. | $\gamma_g^{nl}$ | $4 \cdot 10^{-4}$ | kg/(m<sup>3</sup>.s) | negligible |
| TC metabolism $O_2$ consump. | $\gamma_0^{nl}$ | $6 \cdot 10^{-4}$ | kg/(m<sup>3</sup>.s) | $6.65 \cdot 10^{-4}$ |
| Start TC growth inhibitory | $p_1$ | 1800 | Pa | 1432 |
| Stop TC growth | $p_{crit}$ | 4000 | Pa | 5944 |

### Validation

Unpublished experimental results of encapsulated MCTS, both thick and thin, have been used as validation dataset (Fig. ??E). Each capsule had its own radius *R* and thickness *h* and two simulations have been run with the extreme experimental values of the alginate stiffness (*E*_alg_ = 47 kPa and *E*_alg_ = 89 kPa).

Fig. 6A right shows the range of modeling possibilities of the identified parameters on the training data CCT0, respectively to the alginate stiffness range. The parameters set was identified with the mean stiffness value (*E*_alg_ = 68 kPa).

Fig. 6B shows that the modeling range on two thick capsules, CCT1 and CCT2, is in accordance with the experimental results. Two fits with an alginate stiffness at *E*_alg_ = 52.5 kPa and *E*_alg_ = 70 kPa respectively are proposed.

The Fig. 6C shows results relative to two thin capsules, CCT3 and CCT4. The dynamic is properly reproduced by the model for both capsules which are importantly deformed (the strain is of 16% for the left one and 20% for the right one). Despite in the right case CCT4 where the model shows some limitations, because the proposed fit is at the minimum of the experimental stiffness value *E*_alg_ = 47 kPa, the presented cross-validation demonstrates that this mechanistic mathematical model can adapt to different geometries and thickness without losing its relevance. Focusing on the left graphs in figures B and C we can note that, with the same parameters set and almost the same alginate stiffness (B Left, *E*_alg_ = 52.5 kPa and C Left, *E*_alg_ = 54 kPa), the model reproduces experimental strain of 8% and 16% respectively. The difference between the two strains is induced by the geometrical effect due to the capsule thickness, which impacts on the evolution of internal stresses, cell growth and oxygen consumption.

### Qualitative results and emerging outcomes

In addition to overall quantitative results, Fig. 7 and Fig. 8 provide details on the physical phenomena occurring during growth (from confluence to 85 hours after confluence) of a MTCS encapsulated in a thick capsule with the same geometry than CCT0. These figures quickly allow understanding the importance of physics-based modeling, as it provides qualitative information that could be used to interpret the experimental process as a whole and to better understand the tumor growth process. Fig. 7 shows contours of oxygen, necrotic fraction, IF pressure, ECM displacement, TC pressure and TC saturation at confluence and 85 hours after. To gain information about the dynamics of these quantities, Fig. 8 shows them probed along the radius at confluence, 85 hours after, and at two intermediate times (28 and 56 hours).

**Fig 7.**
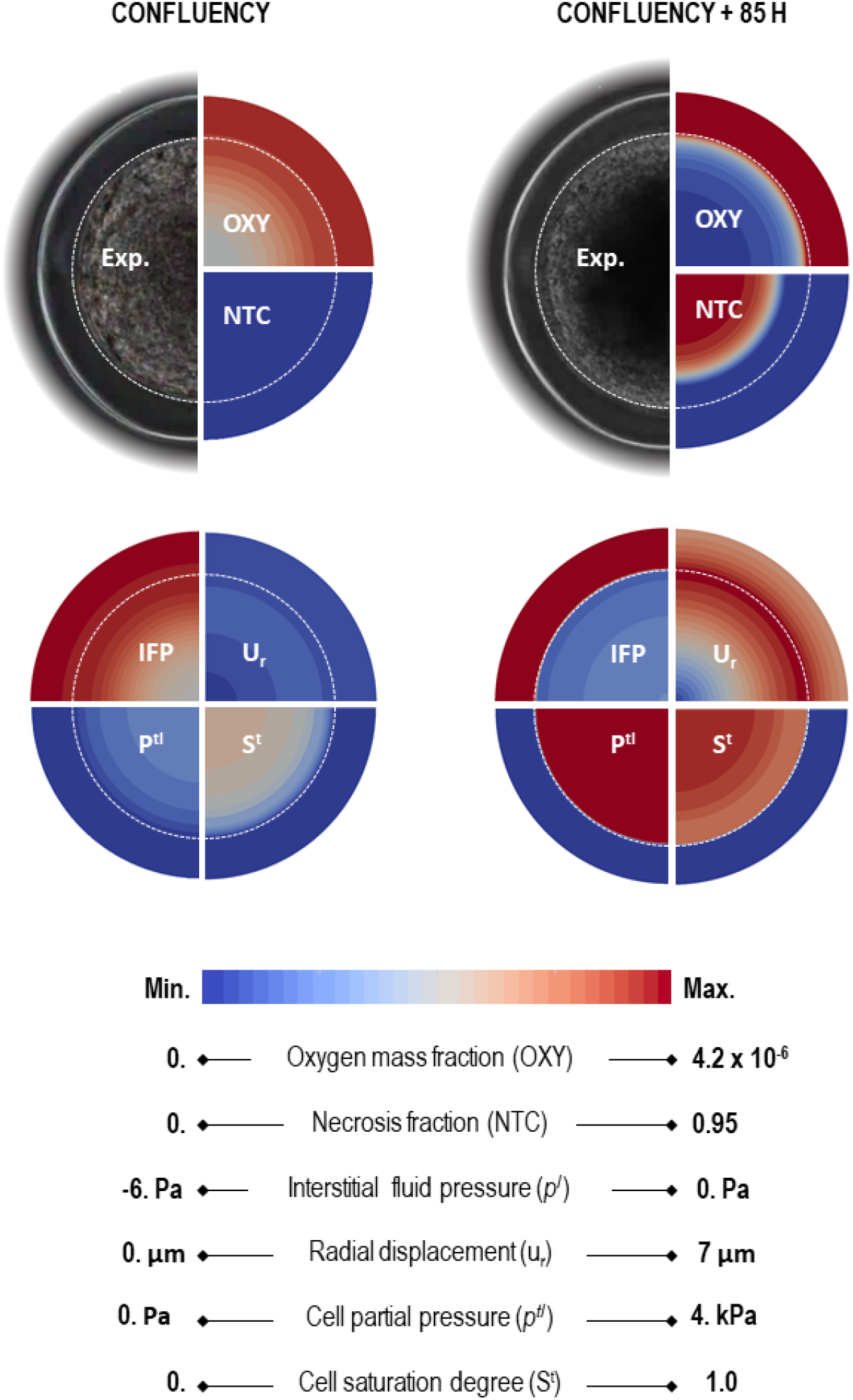
Experimental microscopy image augmented by qualitative numerical results. 6 physical quantities from numerical results of the mathematical model on CCT0: oxygen, necrotic tumor cells, interstitial fluid pressure, radial displacement, the pressure difference between the phases *l* and *t* and tumor cells saturation. Left, at confluence. Right, 85 hours after confluence.

**Fig 8.**
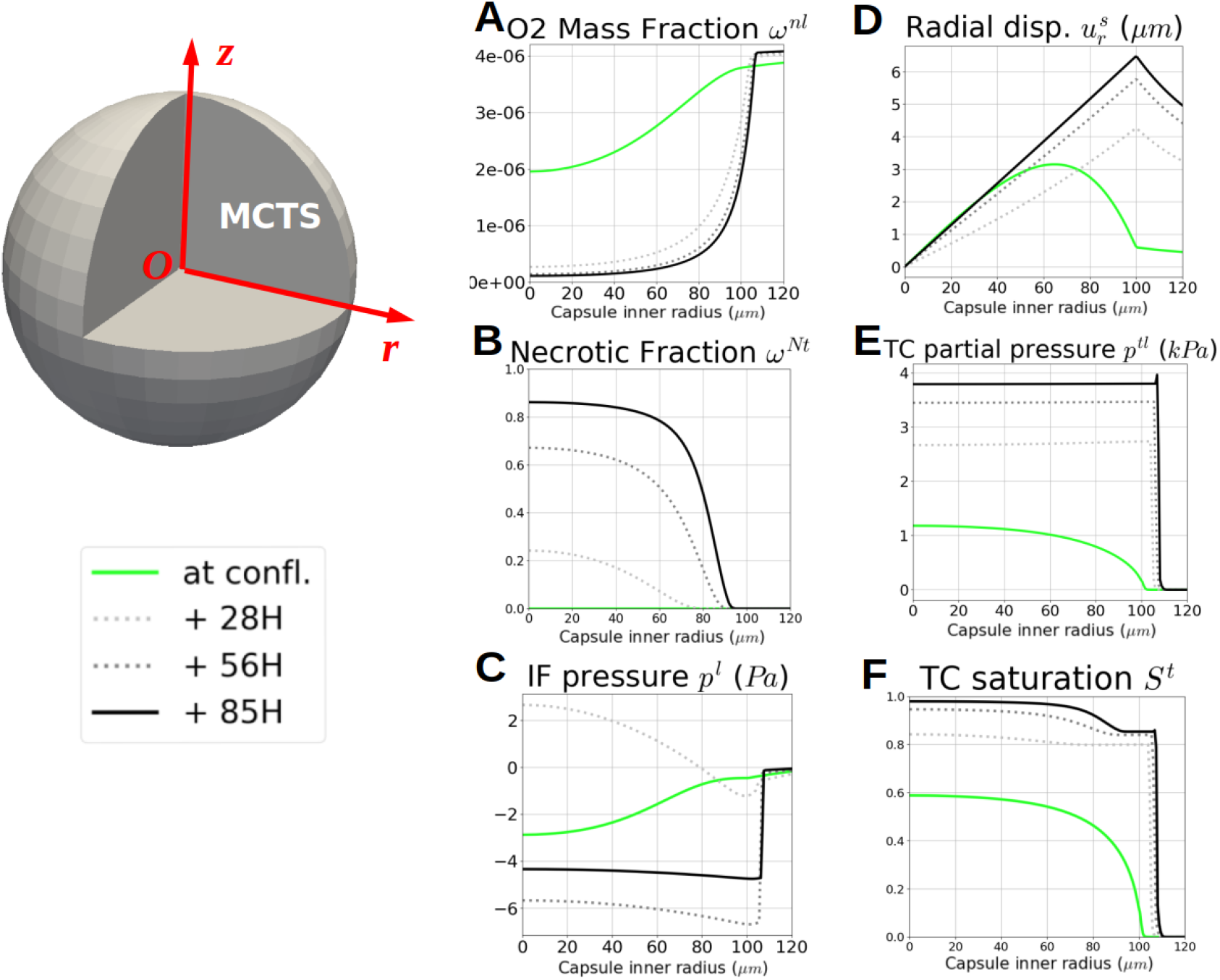
Qualitative numerical results of CCT0 probed along the *r* = *z* line. Quantities probed at confluence, 85 hours after, and two intermediate times (28 and 56 hours): **A** oxygen, **B** necrotic fraction, **C** IF pressure, **D** ECM displacement, **E** TC pressure and **F** TC saturation. **A** and **B**: as mentioned in the experiments, 85 hours after confluence the viable space remaining for TC is a 20 *μ*m thick rim. **C**: after confluence, IF is absorbed by growing TC, provoking a sucking phenomenon, as the cells activity decrease at the MCTS inner core, IF accumulates and its pressure becomes positive. As described in [7], after 2 days of quick growth, the MTCS reaches a state of linear and slow evolution. **D** and **E**: the capsule displacement is driven by TC pressure with the same overall dynamic. **E**, **F** and **B**: relation between the saturation of TC and their necrotic fraction, the TC aggregate density increases with necrotic core.

Fig. 8A and B show the interplay between oxygen consumption and necrosis. Indeed as mentioned in the experiments, 85 hours after confluence, the viable space remaining for TC is a 20 *μ*m thick rim. This is explicit in Fig. 7, upper right circle, NT quarter. The comparison of Fig. 8F and B shows a relation between the saturation of TC and their necrotic fraction. This is a basic experimental fact that, when the cells bodies collapse in a necrotic core, the aggregate density increases accordingly. Fig. 8D and E allow ’visualizing’ the overall dynamics of the process: the capsule displacement strongly increases after confluence due to the contact with tumor cells whose pressure rises from 1.15 kPa at confluence to almost 4 kPa, 85h after confluence. Beyond 85h and until eight days after confluence, it was observed that the tumor cells pressure *p^t^* < *p*_crit_. This is in accordance with the experiment as recorded in [7] where it was reported that the MCTS continued to grow twelve days after confluence, even very slowly.

The tumor cells pressure *p^t^* does not determine directly the capsule deformation, which is more directly driven by the solid pressure, *p^s^* (see definition Eq.21). The pressure *p^s^* is more representative of the average internal pressure Pconf obtained experimentally by inverse analysis (see Eq.3). At the confluence time, *p^s^* is importantly lower than the pressure in the tumor cell phase since at that time the MCTS also consists of 40% of IF. After confluence the saturation of tumor cells increases progressively, so *p^s^*, becomes closer to the pressure sustained by the cells. To allow the numerical comparison, we designed a mesh with a subdomain along the internal radius of the capsule to integrate the numerical quantities with FEniCS. Fig 10 shows the comparison between *p^s^*, *p^t^* and the *P*_conf_.

**Fig 9.**
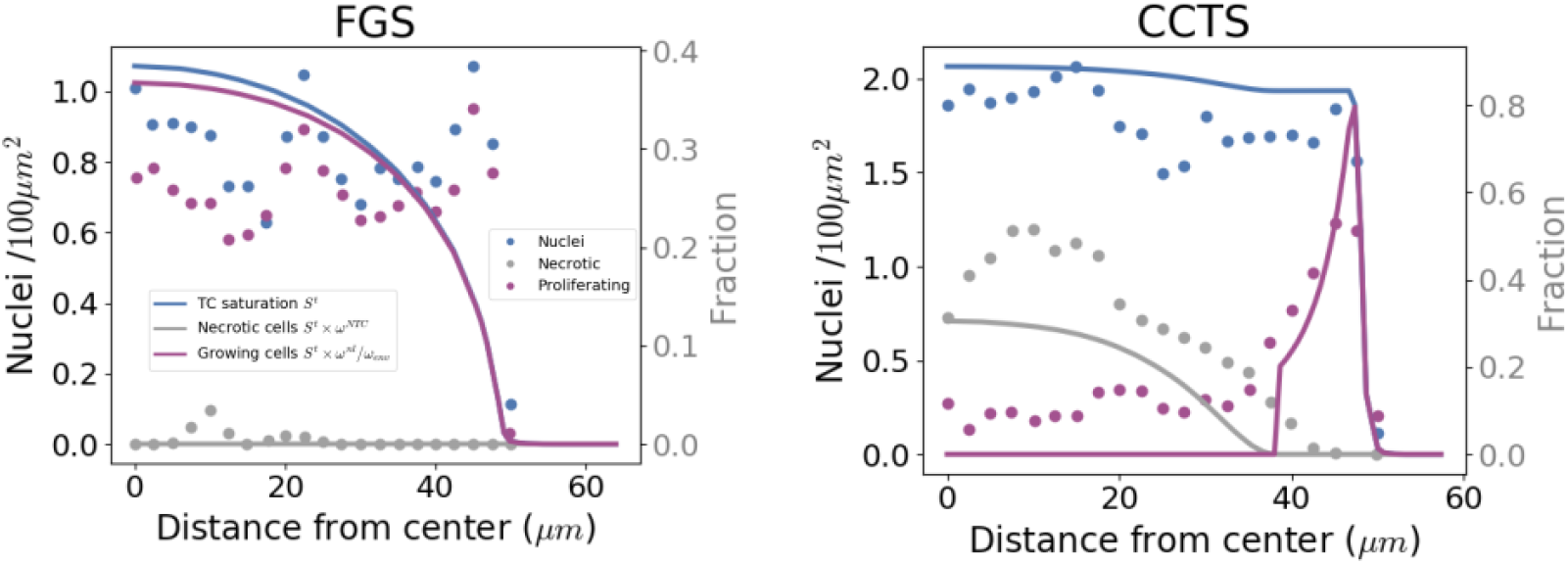
Qualitative comparisons of proliferative and dead cells. Quantification of proliferating and dead cells radial densities for free growth and CCTS: *in vitro-in silico* results. Experimental quantification of cell nuclei (blue), proliferating cells (purple),and dead cells (gray) along the spheroid radius (2. 5 *μ* m sampling). Numerical quantities: TC Saturation *S^t^*, blue dotted; Necrotic saturation of TC *ω^Nt^S^t^*, gray dotted; Growing TC fraction *ω*^*t*^_grow_ (see Eq.31), purple dotted. **A**, FGS. **B**, CCTS, **B**, growth (from [7]). TC saturation almost doubles between the two configurations, in encapsulation, necrotic fraction occupy almost half of the TC phase and only a thin rim of the MTCS is viable. CCTS is very far of the conditions of parameters calibration, nevertheless *in vitro-in silico* comparison shows a reasonably good agreement, which is quantified Table 2.

**Fig 10.**
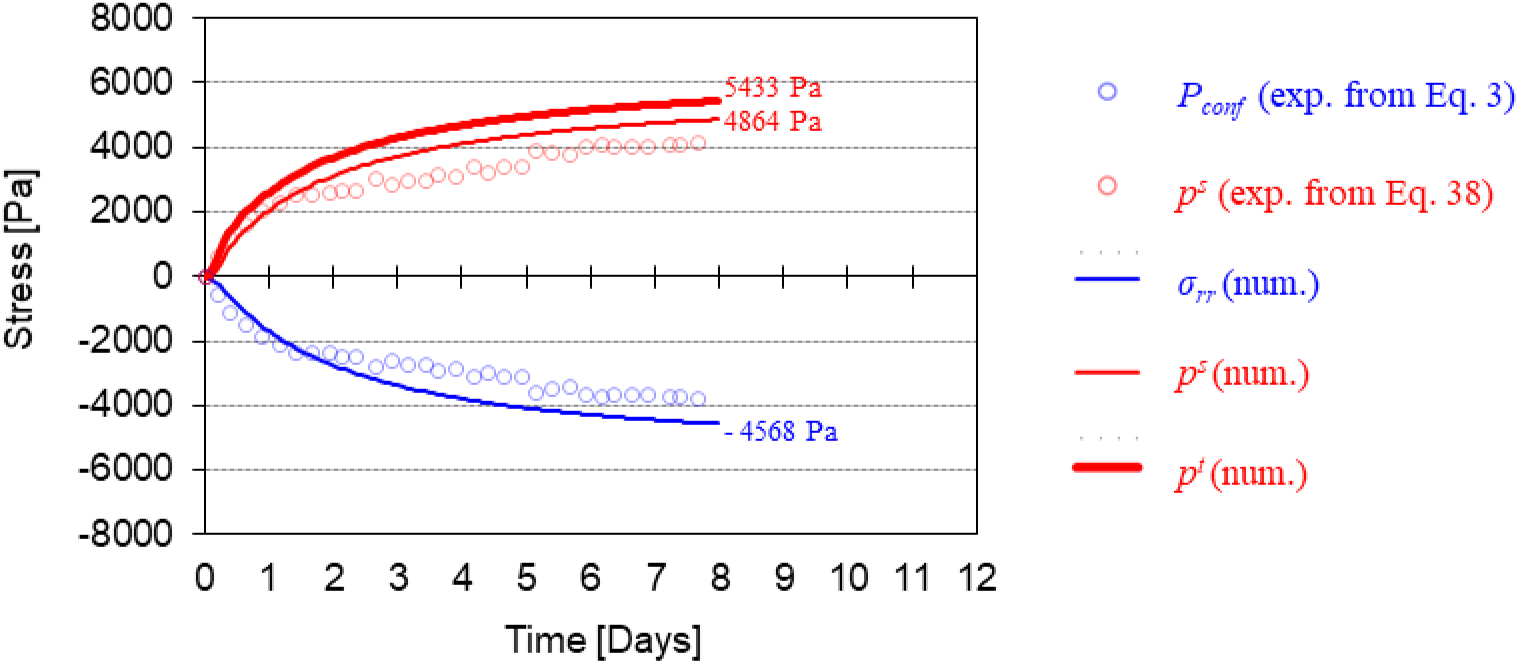
Comparison between numerical stress/pressures and the ones retrieved from the experimental results of CCT0. Blue circles are the confinement pressure obtained experimentally by inverse analysis with equation 3; red circles is the solid pressure, *p^s^*, obtained from the measured displacement using equation 38; solid lines are *σ_rr_, p^s^* and *p^t^* obtained numerically.

In the presented physics-based approach, mass conservation is prescribed, so the growing MCTS, which increases in density and size, results in a decrease in interstitial fluid mass. This result, which cannot be measured experimentally, is shown in Fig. 8 where a sucking phenomenon due to IF absorption by growing TC can be observed. The Fig. 8C shows that after confluence the interstitial fluid pressure becomes positive during a while (see plot relative to 28h). Indeed, after confluence the initial gradient of IF pressure (green line in Fig. 8C) reverses since cells in the proliferative peripheral areas move toward the core so IF has to go in the opposite direction, as imposed physically by mass conservation. After 2 days of quick growth, experimentally and numerically, the MTCS reaches a state of linear and slow evolution and from that point onward, the IF flux will not qualitatively change.

To further analyze the reliability of the mathematical model we also exploit additional data of cell states inside MCTS presented in [7]. More specifically, we reproduced numerically a CT26-MCTS growing in free conditions and the capsule CCTS. Fig. 9A and Fig. 9B present experimental cell densities (total, proliferative and necrotic, plain lines) at 50 *μ*m radius for the free MCTS and 26h after confluence for CCTS. These experimental results are qualitatively compared with the numerical simulations (Fig. 9A and Fig. 9B for the free and confined cases respectively, dotted lines). Both configurations show a reasonable agreement with the experimental results, knowing that none of these quantities have been used for the parameters identification and are very far of the conditions of calibration. This is a supplementary argument that showcases the adaptability of this physical based modeling. The accuracy of the results Fig. 9A and B have been quantified by RMSE (2.5 *μ*m sampling on experimental data) in Table 2. We used the data available in the free MCTS (FGS), to normalize *S^t^* by the experimental nuclei density at *r* = 0. Unfortunately, in this work, we did not have access to a large sample size to evaluate our numerical assumptions against experimental data.

**Table 2.** Quantitative comparison between *in vitro-in silico* results, for FGS radius and CCTS.

| *in vitro-in silico* results | RMSE control group | RMSE CCTS |
| --- | --- | --- |
| nuclei/$S^t$ | 0.317 | 0.486 |
| proliferative/$\omega_{grow}^t S^t$ | 0.303 | 0.327 |
| necrotic/$\omega^{Nt} S^t$ | 0.026 | 0.238 |

### On the experimental estimation of the tumor cell pressure

As explained in the work of Alessandri et al. [7], the pressure in the tumor cell phase is directly related to the confinement pressure arising from the interaction between the capsule and the MCTS calculated with equation 3. However, from the perspective of porous media mechanics, this confinement pressure essentially corresponds to the total Cauchy stress tensor and not to the pressure in the tumor cell phase. Actually, in the post-confluence stage, the rise of cell pressure induces the deformation of the alginate shell but also the deformation of the extracellular matrix (constituting the solid scaffold of the MCTS). Thanks to spherical symmetry of the problem, it is easy to derive an equation which allows us calculating the ’experimental’ solid pressure, *p^s^*, from the confinement pressure (calculted with equation 3) and the stiffness of the ECM (here assumed equal to 1kPa). From the effective stress principle and accounting for assumed elastic constitutive behavior, the solid pressure can be estimated with the following equation

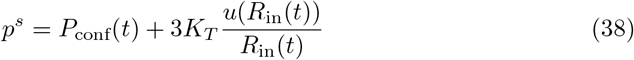

with *K_T_* Bulk’s modulus of the ECM scaffold.

From the solid pressure, the tumor cell pressure could be estimated by means of an experimental evaluation of the tumor cell saturation degree. The experimental confinement stress evaluated with equation 3 and the solid pressure evaluated with equation 38 are depicted in figure 10 for CCT0. In figure 10, numerical results for the temporal evolution of *σ_rr_*, *p^s^* and *p^t^* within the MCTS (proximity of the alginate shell) are also reported. It can be observed that the final tumor cell pressure is almost 20% higher than the magnitude of the computed radial stress. This difference is however strictly related to the Bulk’s modulus of the ECM scaffold for which may also vary during the experiment due to the production of extracellular matrix by tumor cells. The mechanical properties of the ECM have not been investigated in this study, thus these results are theoretically interesting but only have a qualitative value.

A deviation of the solid pressure and the radial stress from respective experimentally-derived values can be observed in the figure. The reason is that equation 3 has been derived assuming a non compressible alginate while in our calculation we assumed a Poisson ratio *ν* = 0.4. When strain becomes relatively large in the numerical case the reduction of the thickness due to geometrical non-linearity is smaller than that of the assumed non compressible case, so the alginate shell is more rigid and consequently the pressure is higher.

## Discussion

We showed, in this paper, the *in silico* reproduction of MCTS growth experiments in various physical conditions: free and encapsulated within alginate shells of different sizes and thicknesses. Thanks to a robust validation protocol, variance-based sensitivity analysis, distinct training and validation datasets, all these physical conditions have been successfully simulated by means of a bio-chemo-poromechanical mathematical model. It is important to notice that only one set of parameters, identified on a training dataset (reported in Fig. 1B and C), has been used for all the numerical simulations performed.

In the frame of the parameter identification process, a local second-order sensitivity analysis has revealed that the parameters of the model become almost independent under confinement (see Fig. 5D) within a range of ±10% around the usual literature values. Results of sensitivity analyses also demonstrate that, if the tumor is free to grow, the only influential parameters are the proliferation and the oxygen consumption rates. Conversely, when the tumor growth is constrained by the presence of the alginate capsule, the value of the critical pressure beyond which mechanical stresses inhibit its growth is the main driver. The mathematical model remains reliable even when the growth conditions of the MCTS are modified. This is an advancement with respect of other numerical studies based on poromechanics which are quite qualitative [23] or solely connected with a reference experimental setup [20].

However, this advancement is only a first step to a more robust *in vitro/in silico* process. From experimental side, a wider set would raise the confidence on the calibration of the parameters and the validation of the model; repeating this experimental protocol on other standardized cancer cell-lines (glioma, breast cancer, prostate cancer, to name a few) would allow enlarging the understanding if this mechano-biological framework and its digital twinning. From the numerical side, if a detailed but local sensitivity analysis accelerate the calibration process, its relevance concerns only a part of the parameters’ space, and only these experimental data. Attempt of global sensitivity without a data bias, or extent these local analyses to other cell-line would be a desirable next step.

The mathematical model is the digital twin MCTS-capsule system since it takes into account mechanistically its real multiphase nature; hence, the numerical results add new dimensions to the Cellular Capsule Technology. In particular, it is shown that the pressure estimated experimentally is illustrative of the evolution of the solid pressure, *p^s^*, (in the sense of porous media mechanics, see Eq.21) and not of the pressure sustained by the cells, *p^t^*. The pressure *p^t^* is always higher than *p^s^* especially during the first phase after confluence (when the MTCS still contains an important volume fraction of IF). This fact is the direct consequence of the fact that each phase of the MCTS (*i.e.,* the ECM, the IF and the TC) has its own stress tensor and that the pressure obtained experimentally by inverse analysis is an average pressure (see Fig. 10). The multiphase approach also reveals other behaviors not measurable experimentally. We observe for instance that after confluence there is a suction of IF from the extra-capsular domain and that cells move from the proliferating rim towards the core of the MCTS where they become necrotic.

Although these emerging results are inspiring, several physical phenomena are not represented and could lead to valuable insight from the experimental and modeling point of view. The mechanical stress can be the primer of cell necrosis [3] and the CCT experimental framework would be a interesting framework to measure this mechano-biological interplay. The phenotype switch of tumor cells under homeostatic pressure studied in [2] could be modeled and revealed by a alternating CCT growth and free growth on the same aggregate. In this specific study, we hold a line which we wanted simple, by limiting the number the modeled physical phenomena, light, our code can be run on a eight-core processor, and easily adaptable to any geometry and cell-line.

In 2020, mathematical modeling in oncology begins to enter a stage of maturity; today mathematical models of tumor growth tend to clinical applications and therefore must be really predictive and funded on measurable or at least quantifiable parameters having, as much as possible, a sound physical meaning. This motivated this paper which presents not only a mechanistic bio-chemo-poromechanical model but also a *modus procedendi* to achieve a suitable predicative potential and, with intercession of sensitivity analysis, to quantify relative relevance of mechanisms underlying tumor growth phenomenology.

## Acknowledgments

The results presented in this paper were carried out using the HPC facilities of the University of Luxembourg [31] (see https://hpc.uni.lu).

## Appendix A. Computational framework

The model has been coded in Python and C++ in the open-source FEniCS framework [30] with an incremental monolithic resolution of the mixed finite element (FE) formulation. The monolithic resolution allows us reducing substantially the computational time compared with staggered resolution methods usually adopted (*e.g.,* see [19]). Whereas spherical symmetry is assumed in experimental results, we have chosen cylindrical symmetry to preserve the generality and the adaptability of the FE mesh and formulation. Even if the computational time is more important, it remains reasonable: 3 hours in a single core of an average laptop; 1D spherical formulation would have forced us to quit classical FE formulation or to design, for each case, a specific finite difference formulation.

An updated lagrangian approach has been adopted to account for geometrical nonlinearities, the incremental resolution allows us updating primary variables as follows:

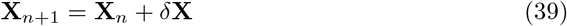

with *δ***X** the vector of unknowns

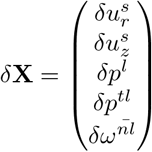

After each time step, the space *χ*^*s*^ ∈ ℝ^2^ is updated:

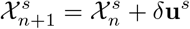

### Choice of the element

For all mixed FE problem with vectorial and scalar coupled unknowns, the chosen finite element should verify the inf-sup condition, that is to say, should preserve the coercivity of the bilinear form (see [32] p.223-230). A simple choice is the Taylor-Hood element, with a Lagrange element of order *k* ≥ 1 for the scalar unknowns and order *k* + 1 for the vectorial one. However, modelling an encapsulated tumor growth implies a very sharp gradient at the capsule inner radius for the pressure difference *p^tl^*, between *l* and *t* phases. The linear approximation of the Lagrange element of order 1 could not describe it, except at the cost of an extremely refined mesh at the interface, and the error could provoke *numerical infiltration* of tumor cells in the alginate capsule (see Fig. 11). To avoid this phenomenon, the composite Taylor-Hood element has been set to a higher order, precisely the mixed FE formulation in FEniCS uses the composite Taylor-Hood element *P*_3_(ℝ^2^), [*P*_2_(ℝ)]^3^. The demonstration of Lax-Milgram theorem for this type of mixed problem could be found in the *Encyclopedia of Computational Mechanics,* Vol.1, p.149-202 [33].

**Fig 11.**
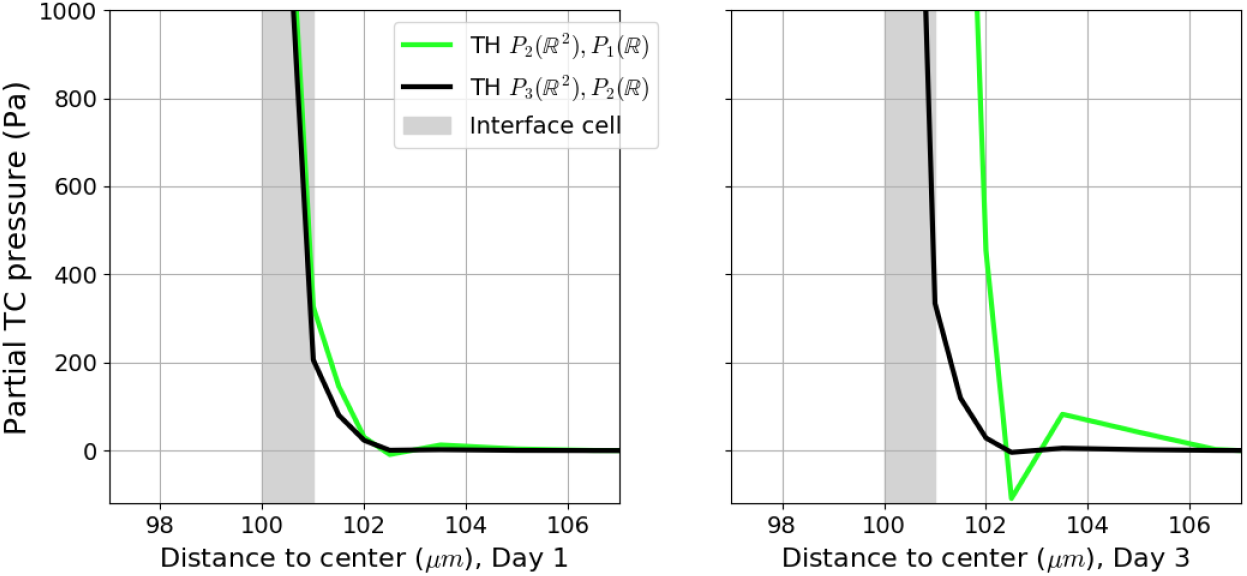
Choice of the element. Composite Taylor-Hood *P*_2_(ℝ^2^), [*P*_1_(ℝ)]^3^, green; composite Taylor-Hood *P*_3_(ℝ^2^), [*P*_2_(ℝ)]^3^, black. The linear approximation *P*_1_(ℝ) of the pressure difference between *l* and *t* phases at the capsule interface (Interface element shared, green) is poor (Left, Day 1) and provoke numerical infiltration of tumor cells into the alginate capsule (Right, Day 3). The quadratic approximation *P*_2_(ℝ) (Interface element shared, black) does not provoke numerical infiltration.

### Choice of mesh cell size

The mixed FE problem has been computed on 5 different meshes, with uniform cell sizes *dh* = 50,20,10, 5 and 2.5 *μ*m. To measure the FE solution degradation the primary variable 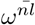, the oxygen mass fraction, has been monitored at the spheroid center for 4 days (see Fig Fig. 12). The thinner mesh of cell size *dh* = 2.5 *μ*m has been used as a reference for the RMSE. Despite an important increase of the computation time, the mesh cell size of *dh* = 5 *μ*m has been chosen to restrict the relative degradation of the FE solution to *RMSE* = 0.01 (see Table 3).

**Fig 12.**
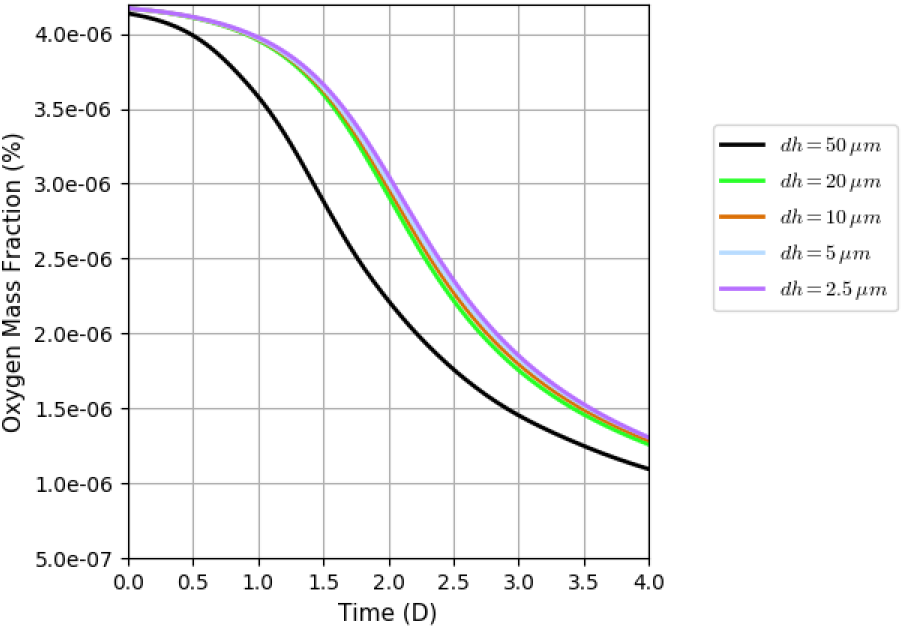
Sensitivity of the solution related to mesh refinement. The numerical oxygen mass fraction, 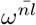, at the center of the spheroid has been monitored during six days for five different mesh finite element sizes *dh*. (*dh* = 50 *μ*m, black; *dh* = 20 *μ*m, green; *dh* =10 *μ*m, brown; *dh* = 5 *μ*m, light blue; *dh* = 2.5 *μ*m, purple)

**Fig 13.**
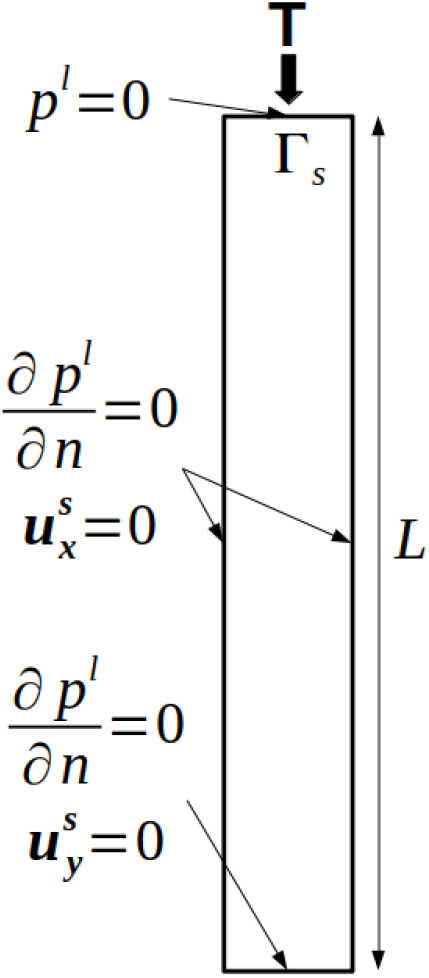
Boundary conditions of the Terzaghi’s problem. The load is applied at the top face, where the fluid is free to espace, say under drained condition. The five remaining faces are under slip condition for the displacement **u**^*s*^ and no flux condition for the fluid pressure *p^l^*.

**Fig 14.**
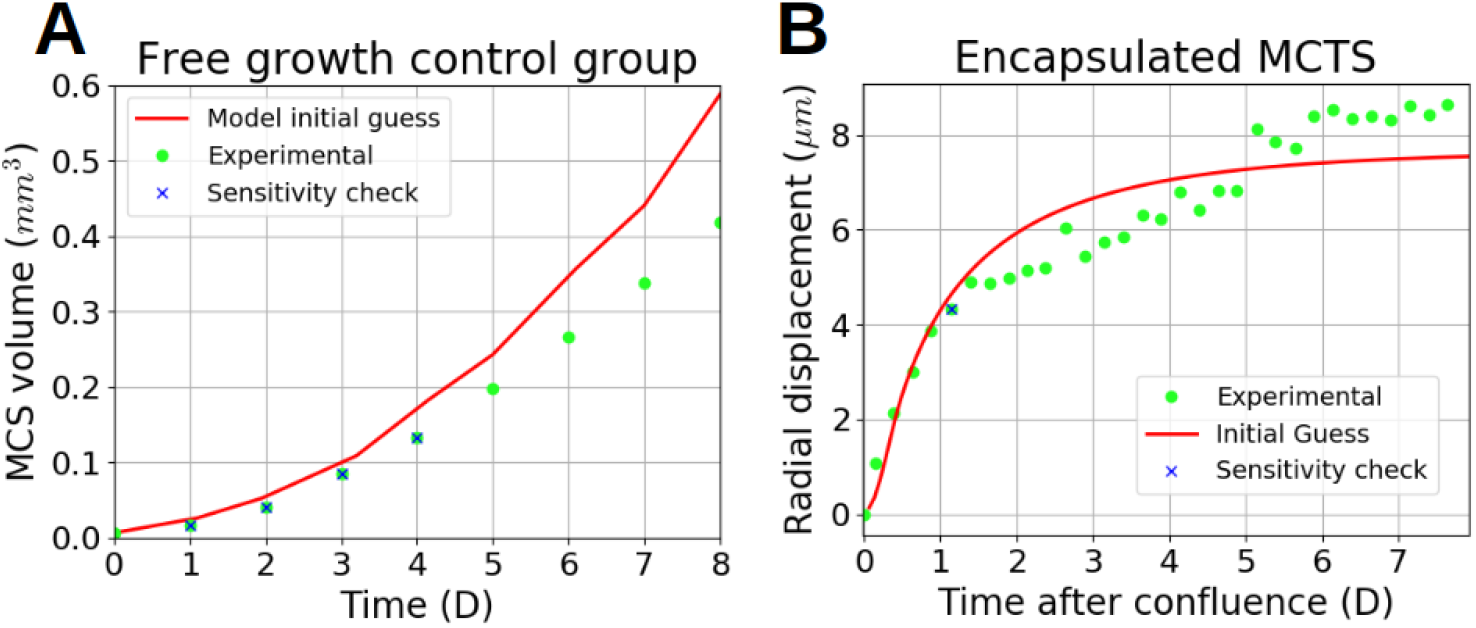
Results with the 7 parameters at their initial values. 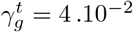, 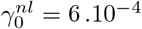, *a* = 800, *p*_1_ = 1800, *p*_crit_ = 4000 (see Table 1). Experimental, green dot; Model, red; Sensitivity evaluation, blue x. Left: FG0; Right: CCT0

**Table 3.**
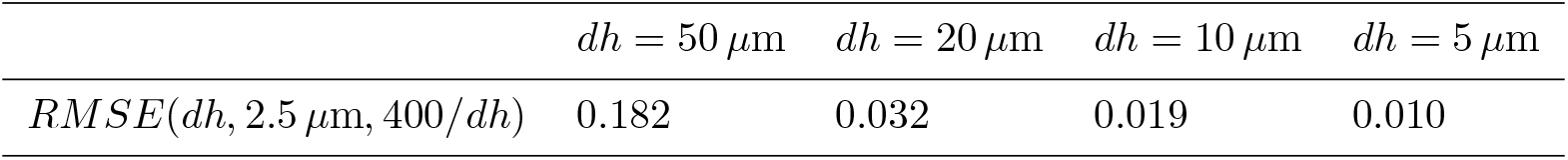
Relative degradation of the solution due to mesh cell size. Measured by root mean square, the reference being the thinner mesh with a mesh cell size of *dh* = 2.5 *μ*m.

| | $dh = 50 \mu\text{m}$ | $dh = 20 \mu\text{m}$ | $dh = 10 \mu\text{m}$ | $dh = 5 \mu\text{m}$ |
| --- | --- | --- | --- | --- |
| $RMSE(dh, 2.5 \mu\text{m}, 400/dh)$ | 0.182 | 0.032 | 0.019 | 0.010 |

### Verification of the FE formulation with an analytical solution

If this system is considered with a single-phase flow into a porous medium under a constant load with the right boundary conditions, one obtains the problem as known as Terzaghi’s consolidation, which has an analytical solution [34]. The system, under a constant load **T**, is reduced to two primary variables the displacement of the solid scaffold **u**^s^ and the pressure of the single phase fluid *p^l^*:

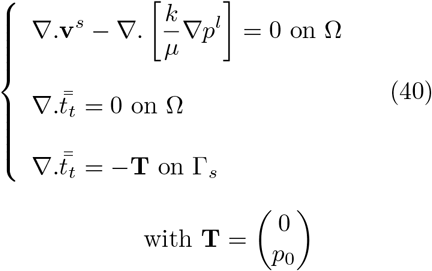

The fluid is free to escape only at the loaded boundary, this boundary condition is known as drained condition. The analytical solution of this problem is:

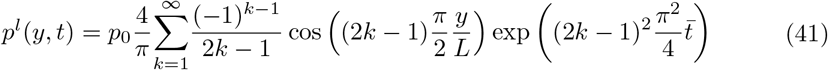

With the characteristic time of the consolidation 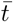, equal to 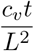, *L* sets to 100 *μ*m and *c_v_*, the consolidation coefficient:

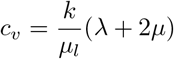

where λ and *μ* are Lamé constants of the solid scaffold, *k* is its intrinsic permeability and *μ_l_* the fluid dynamic viscosity. The addition of the RMSE of the 4 samples at 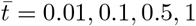 (see Fig. 15) with the analytical solution as reference gives Σ *RMSE* = 0.0028. The surface error for different cell sizes *dh* and time steps *dt* is in Fig. 15(right).

**Fig 15.**
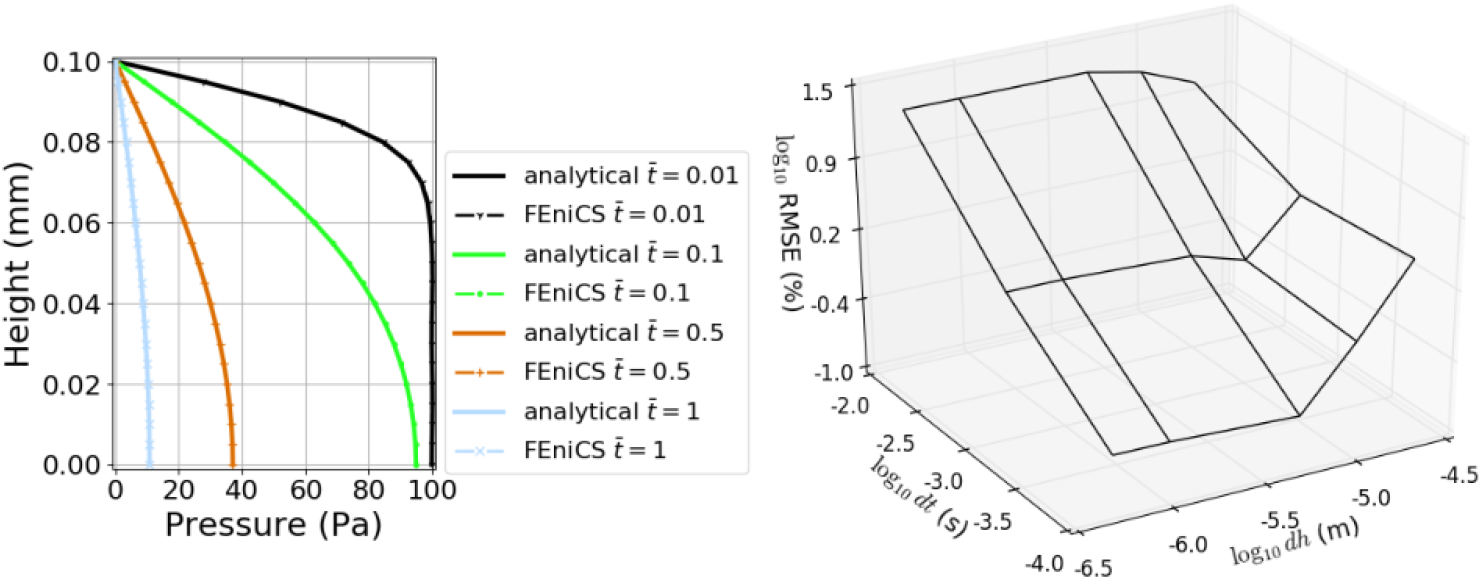
Comparison between analytical solution and numerical result. Left: qualitative comparison between analytical solution of Terzaghi’s problem and FEniCS computation. (4 comparisons at characteristic time of consolidation 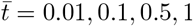). Right: quantitative comparison, error surface between Terzaghi’s analytical solution and FEniCS computation. (x axis: log_10_ of cell size *dh*; y axis: log_10_ of *dt*; z axis: log_10_ of RMSE). The minimum RMSE= 0.0028 is reached at *dh* = 5*μ*m, *dt* = 1*e*^−4^

## Appendix B. Sensitivity analysis

For the sensitivity analysis, the experimental input data were:

- for the free MCTS, the volume monitored over a time span from day 1 to day 4. These data are denoted 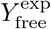
- for the encapsulated MCTS CCT0 the capsule strain one day after confluence and the corresponding analytical pressure (i.e. incompressible elastic membrane). We chose the capsule of inner radius = 100 *μ*m and thickness = 34 *μ*m, presented as the reference case in [7]. These data are denoted 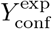

We performed a variance-based sensitivity study of the FE solution on the parameters, both on the free and encapsulated MCTS, as follows:

- A first-order analysis, the 7 parameters are disturbed one at a time respectively to an 8-points grid.
- Interaction analysis, the 21 parameters tuples are evaluated at the 2 extreme points of the grid.

All the results were interpreted with a polynomial model in order to quantify their weights in the FE solution variance, referred to as Sobol indices.

### First-order analysis

Each parameter is disturbed one at a time respectively to this grid [–10, –5, –2, –1, +1, +2, +5, +10]%, giving the corresponding 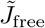 and 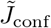. The relative variations of the cost functions were calculated as follows:

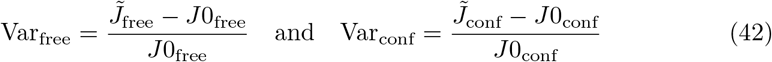

where *J*0_free_ and *J*0_conf_ are the costs with the parameters at their generic values. In order to quantify the impact of each parameter, the following linear model was set:

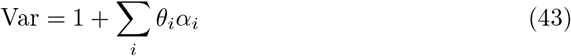

where *α_i_* is an auxiliary parameter ∈ [−1, +1] representing the perturbations of the *i^th^* parameter along the grid and *θ_i_* the slope of the variation.

In a first-order analysis, the influence of the *i^th^* parameter is given by the Sobol indices:

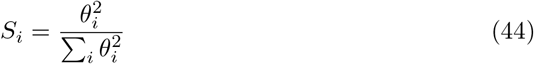

The results for the free and encapsulated configurations are reported in Tables 4 and 5.

**Table 4.** Sobol indices of the first-order sensitivity analysis of the FG0 configuration.

| Parameter | $\theta$ | $S_i(\%)$ |
| --- | --- | --- |
| $a$ | 0.2554 | 6.25 |
| $\mu_t$ | -0.1998 | 3.82 |
| $\gamma_g^t$ | 0.9205 | 81.17 |
| $\gamma_g^{nl}$ | -0.1161 | 1.29 |
| $\gamma_0^{nl}$ | -0.2790 | 7.45 |
| $p_1$ | 0 | 0 |
| $p_{crit}$ | 0 | 0 |

**Table 5.**
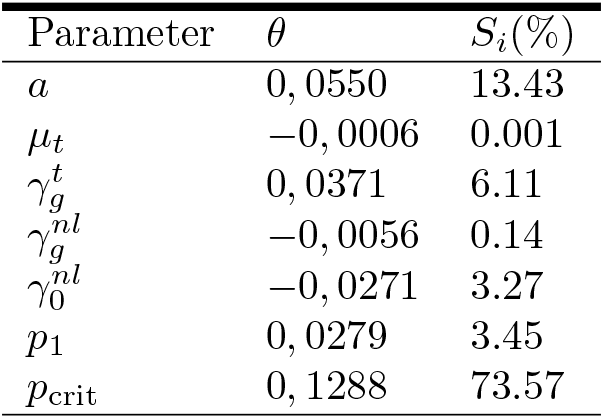
Sobol indices of the first-order sensitivity analysis of the encapsulated growth configuration CCT0.

| Parameter | $\theta$ | $S_i(\%)$ |
| --- | --- | --- |
| $a$ | 0,0550 | 13.43 |
| $\mu_t$ | -0,0006 | 0.001 |
| $\gamma_g^t$ | 0,0371 | 6.11 |
| $\gamma_g^{nl}$ | -0,0056 | 0.14 |
| $\gamma_0^{nl}$ | -0,0271 | 3.27 |
| $p_1$ | 0,0279 | 3.45 |
| $p_{crit}$ | 0,1288 | 73.57 |

### Interaction analysis

As the independence of physical phenomenons involved in encapsulated configuration is one our major modeling assessment, the interaction between parameters has also been studied. The 21 tuples have been evaluated at the 2 extreme values of the grid for each configuration. The corresponding polynomial model becomes:

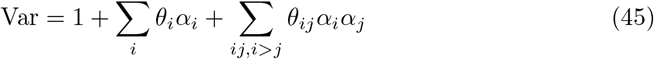

with the respective Sobol indices:

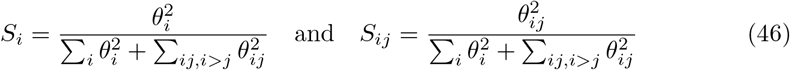

The results for the free and encapsulated configurations are reported in Tables 6 and 7.

**Table 6.**
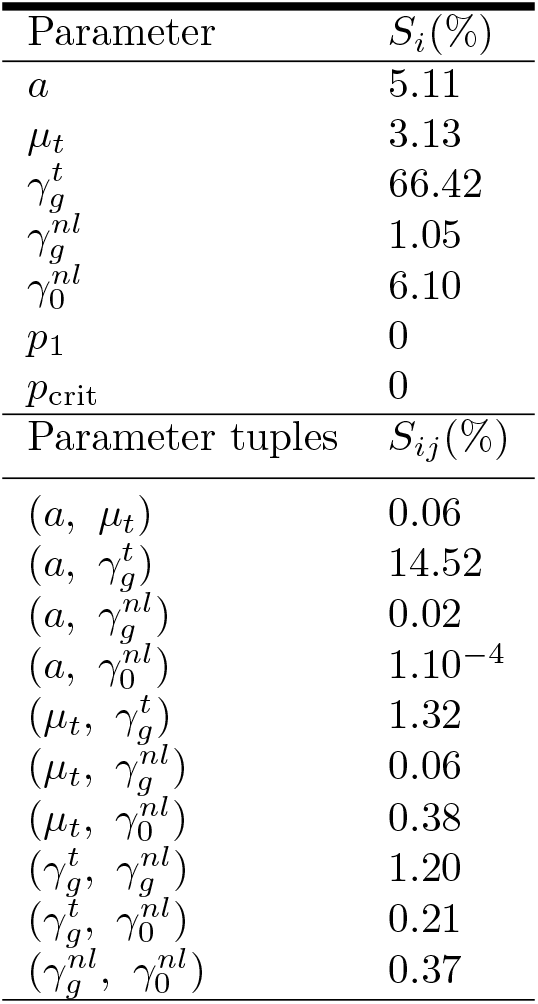
Sobol indices of the interaction sensitivity analysis of the FG0 configuration.

**Table 7.** Sobol indices of the interaction sensitivity analysis of the encapsulated growth configuration CCT0.

| Parameter | $S_i(\%)$ |
| --- | --- |
| $a$ | 12.96 |
| $\mu_t$ | 0.001 |
| $\gamma_g^t$ | 5.89 |
| $\gamma_g^{nl}$ | 0.13 |
| $\gamma_0^{nl}$ | 3.15 |
| $p_1$ | 3.33 |
| $p_{\text{crit}}$ | 70.94 |
| Parameter tuples | $S_{ij}(\%)$ |
| $(a, \mu_t)$ | 0.01 |
| $(a, \gamma_g^t)$ | 0.003 |
| $(a, \gamma_g^{nl})$ | 0.01 |
| $(a, \gamma_0^{nl})$ | 0.02 |
| $(a, p_1)$ | 0.009 |
| $(a, p_{\text{crit}})$ | 0.02 |
| $(\mu_t, \gamma_g^t)$ | 0.02 |
| $(\mu_t, \gamma_g^{nl})$ | $5.10^{-6}$ |
| $(\mu_t, \gamma_0^{nl})$ | 0.007 |
| $(\mu_t, p_1)$ | 0.003 |
| $(\mu_t, p_{\text{crit}})$ | 0.7 |
| $(\gamma_g^t, \gamma_g^{nl})$ | 0.02 |
| $(\gamma_g^t, \gamma_0^{nl})$ | 0.01 |
| $(\gamma_g^t, p_1)$ | $5.10^{-4}$ |
| $(\gamma_g^t, p_{\text{crit}})$ | 0.08 |
| $(\gamma_g^{nl}, \gamma_0^{nl})$ | 0.01 |
| $(\gamma_g^{nl}, p_1)$ | 0.002 |
| $(\gamma_g^{nl}, p_{\text{crit}})$ | 0.87 |
| $(\gamma_0^{nl}, p_1)$ | 0.01 |
| $(\gamma_0^{nl}, p_{\text{crit}})$ | 1.30 |
| $(p_1, p_{\text{crit}})$ | 0.42 |

